# Unique recognition, phagocytosis, and intracellular survival of *Cryptococcus neoformans* in a human immortalized microglia cell line

**DOI:** 10.1101/2025.07.17.665466

**Authors:** Robbi L. Ross, Kassandra Arias-Parbul, Zane M. Douglass, Katrina L. Adams, Felipe H. Santiago-Tirado

## Abstract

*Cryptococcus neoformans,* the etiological agent of cryptococcal meningitis (CM), is a globally distributed environmental yeast that mainly causes infections in immunocompromised individuals. Particularly in low-resource countries, the mortality rate of CM can reach 81% and accounts for 19% of HIV/AIDS-related deaths each year. In immunocompromised individuals, once inhaled, *C. neoformans* escapes from the lungs and disseminates with special predilection for the central nervous system (CNS). Once in the brain, *C. neoformans* interacts with microglia, the tissue-resident macrophages of the CNS. Previous studies indirectly showed that microglia are ineffective at controlling this fungal infection. The mechanisms underlying this fungal survival and proliferation within the CNS, however, remain unclear. In this study, we use and validate the C20 immortalized human microglia cell line to study cryptococcal-microglia interactions. We show that microglia have limited phagocytic activity that is specific to *C. neoformans* and partly dependent on cryptococcal antiphagocytic proteins that alter cell size and cell wall structure. We also show human microglia respond to cryptococcal strains differently than peripheral macrophages. Further, we show that human microglia are ineffective at killing phagocytosed *C. neoformans*, and that this could be due to the ability of this yeast to disrupt phagosome maturation and induce phagosome membrane damage in these cells. These findings provide us with fundamental knowledge regarding cryptococcal pathogenesis in the CNS, specifically insight into how *C. neoformans* is recognized by microglia under different conditions and demonstrate the usefulness of C20 cells to further study how this yeast survives and replicates within the CNS environment.

**Importance:** While *Cryptococcus neoformans* is acquired through inhalation, the fatal pathology of cryptococcal infection occurs when the yeast disseminates to the central nervous system (CNS) and causes cryptococcal meningitis. Microglia are the first immune cells that *C. neoformans* will encounter once it reaches the CNS, and they are the largest population of macrophages in the brain. While microglia are professional phagocytes, they are unable to control *C. neoformans* infection. The mechanisms behind uncontrolled growth of *C. neoformans* within the CNS remains understudied, partly due to a lack of knowledge regarding microglia-cryptococcal interactions. This study provides fundamental knowledge into these interactions and establishes a powerful model to specifically study how *C. neoformans* is recognized by microglia and how cryptococcal phagosomes mature in these phagocytes. This work opens new avenues of research to further our understanding of cryptococcal-host interactions which can be leveraged to develop more effective therapeutics for cryptococcal meningitis.

## 1. Introduction

Of the four to five million fungal species worldwide (1), few are frequent pathogens of humans (2). However, this small number of fungi causes over 80 million invasive infections annually, resulting in 3.8 million deaths (3). One of the most common and challenging infections is caused by the environmental yeast *Cryptococcus neoformans* (4). *C. neoformans* is globally distributed, hence we are all frequently exposed, and is responsible for about 147,000 annual deaths, mostly in immunocompromised individuals (3, 5). Desiccated yeasts and spores are inhaled and become lodged in the lungs. In immunocompetent individuals, the lung-resident macrophages (alveolar macrophages, AMs) are able to control or clear the infection. However, under immunocompromised conditions, the yeast disseminates from the lungs with special predilection for the central nervous system (CNS), resulting in lethal meningitis or meningoencephalitis (commonly known simply as cryptococcal meningitis). Cryptococcal meningitis is a leading cause of death in patients with HIV/AIDS and has a general mortality rate of 76% (3, 5). The interactions of immune cells, particularly tissue-resident macrophages, with this fungus are critical determinants of the disease outcome (6). Tissue-resident macrophages are in every organ and function in maintaining homeostasis and immune surveillance to provide rapid defense against invading pathogens. Within the CNS, there are several tissue-resident macrophages present in different anatomical locations. The most numerous and well-studied of these include microglia, which are in the brain parenchyma (7).

Microglia have several functions as the brain’s resident immune cells, including but not limited to synapse formation and degradation, engulfment of apoptotic neurons and neurotransmitters, surveillance of neuronal activity, and rapid responses to tissue damage and invading pathogens. Several pathogens, including bacteria, viruses, fungi, and parasites, are able to invade and cause infection within the CNS. In most cases, microglia are the first immune cells these pathogens will encounter once they cross the blood-brain barrier (BBB) and have been shown to be critical in controlling several of these pathogens. For example, microglia rapidly respond to the parasite *Trypanosoma brucei*, leading to its engulfment and elimination (8). Furthermore, the expression of Toll-like receptor 4 (TLR-4) and CD11b on microglia is responsible for the clearance of *Candida albicans* from the brain (9).

While the lethal pathology of cryptococcal infection occurs in the brain, the interactions of *C. neoformans* with microglia remain understudied. However, it has been shown that microglia play a permissive, rather than protective, role during cryptococcal infection. *In-vivo* experiments show rapid expansion of *C. neoformans* in the brain does not immediately activate murine microglia nor cause them to upregulate MHC class II expression (10, 11). Additionally, *ex-vivo* and *in-vivo* histological analyses show that *C. neoformans* survives and replicates within human microglia, implying limited microglia fungicidal activity (12–14). The mechanisms responsible for this uncontrolled growth of *C. neoformans* and lack of microglial activation remain unclear, mostly because of a lack of physiologically relevant *in-vitro* models that would allow for mechanistic studies.

In most instances, after recognition and phagocytosis by microglia, a pathogen will reside in a phagosome that undergoes a series of maturation steps becoming a microbicidal lysosome, contributing to control of the infection. However, indirect evidence implies that *C. neoformans* resides in the microglial phagolysosome and resumes intracellular replication (15). In peripheral macrophages, *C. neoformans* manipulates phagosome maturation by altering phagosome acidification, inducing membrane damage, and impairing fungicidal activities (16–23). Despite this, studies extending these findings to microglia are lacking. Furthermore, the similarities and differences regarding cryptococcal interactions between microglia and peripheral macrophages remains understudied. Overall, the factors that influence recognition and engulfment of *C. neoformans* by microglia, and how these subsequently alter phagosome maturation and fungicidal activity, are not well studied.

We set out to study the interactions between *C. neoformans* and microglia before and after engulfment and define how these interactions compare with those observed in peripheral macrophages. We predominately studied these interactions using the immortalized human microglia cell line C20, which has recently been developed and used to study a variety of microglia-HIV interactions as well as microglia cellular processes during neuroinflammation (24–28). Our findings show that the C20 cell line is comparable to primary human microglia and is a useful tool for studying cryptococcal-microglia interactions. Furthermore, we show that *C. neoformans* evades phagocytosis by microglia, partly due to antiphagocytic cryptococcal proteins, in a manner independent of that shown in other macrophage-cryptococcal studies. Additionally, we show that the ability of *C. neoformans* to be engulfed at higher rates by microglia could be related to cell size and cell wall modifications, subsequently altering microglial phagosome maturation. We also show that the ability of *C. neoformans* to survive within human microglia could be due to delayed phagosome maturation and fungal-induced phagosomal membrane damage. Overall, we demonstrate the utility of C20 cells to study the cryptococcal-microglia interactions, allowing for many new lines of investigation into this important topic.

## 2. Results

### 2.1. Microglia have limited phagocytic and fungicidal activity that is specific to *C. neoformans*

To begin studying cryptococcal-microglia interactions, we looked at the ability of C20 cells, primary human microglia, and THP-1 alveolar macrophages (AMs), to engulf the wild-type (WT) *C. neoformans* strain, KN99α (Fig. 1A). The phagocytic index (PI), which represents the number of internalized fungi per 100 macrophages, is comparable between C20 cells and primary human microglia, being 2.3 and 1.7, respectively. Notably, both types of human microglia have a significantly lower PI compared to that of AMs, which is 37. To determine if this lack of phagocytic activity seen with the human microglia is an inherent difference between microglia and AMs, we assessed the ability of C20 cells to engulf BY4741, a common laboratory *Saccharomyces cerevisiae* strain that is used as a control due to its non-pathogenicity and similar cell size and cell wall morphology to *C. neoformans* (Fig. 1B). Interestingly, C20 cells have a significantly higher PI regarding BY4741 co-incubation (average PI = 132.6) than that of KN99α (average PI = 2.56). This indicates that the lack of phagocytic activity from the human microglia for *C. neoformans* (Fig. 1A) is fungal-dependent and not microglia specific. We also saw the same result with primary murine microglia, where they engulf BY4741 to a significantly higher degree than KN99α (average PIs of 119.2 and 44.9, respectively; Fig. 1C). We then tested the ability of C20 cells to inhibit the growth of internalized KN99α and BY4741 (Fig. 1D). We saw that microglia were not effective at controlling the growth of internalized KN99α at 24- or 48-hours after all extracellular fungi were removed. Specifically, after 24-hours, there were 12.74 times more KN99α CFUs than initially engulfed, and this increased to 68 times after 48-hours. However, there was a decrease in BY4741 CFUs after 24- and 48-hours, with there being 0.76- and 0.44- times less fungi than initially engulfed, respectively. This indicates that C20 cells allow for a permissive environment that promotes the intracellular survival and replication of *C. neoformans* whereas, in contrast, have fungicidal activity against BY4741. Overall, this indicates that both C20 cells and primary human microglia have the same poor phagocytic activity for WT *C. neoformans* (KN99α), and these cells might recognize *C. neoformans* differently than AMs. Furthermore, the decreased phagocytic and fungicidal activity seen with microglia seems to be specific for *C. neoformans*.

**Fig. 1.**
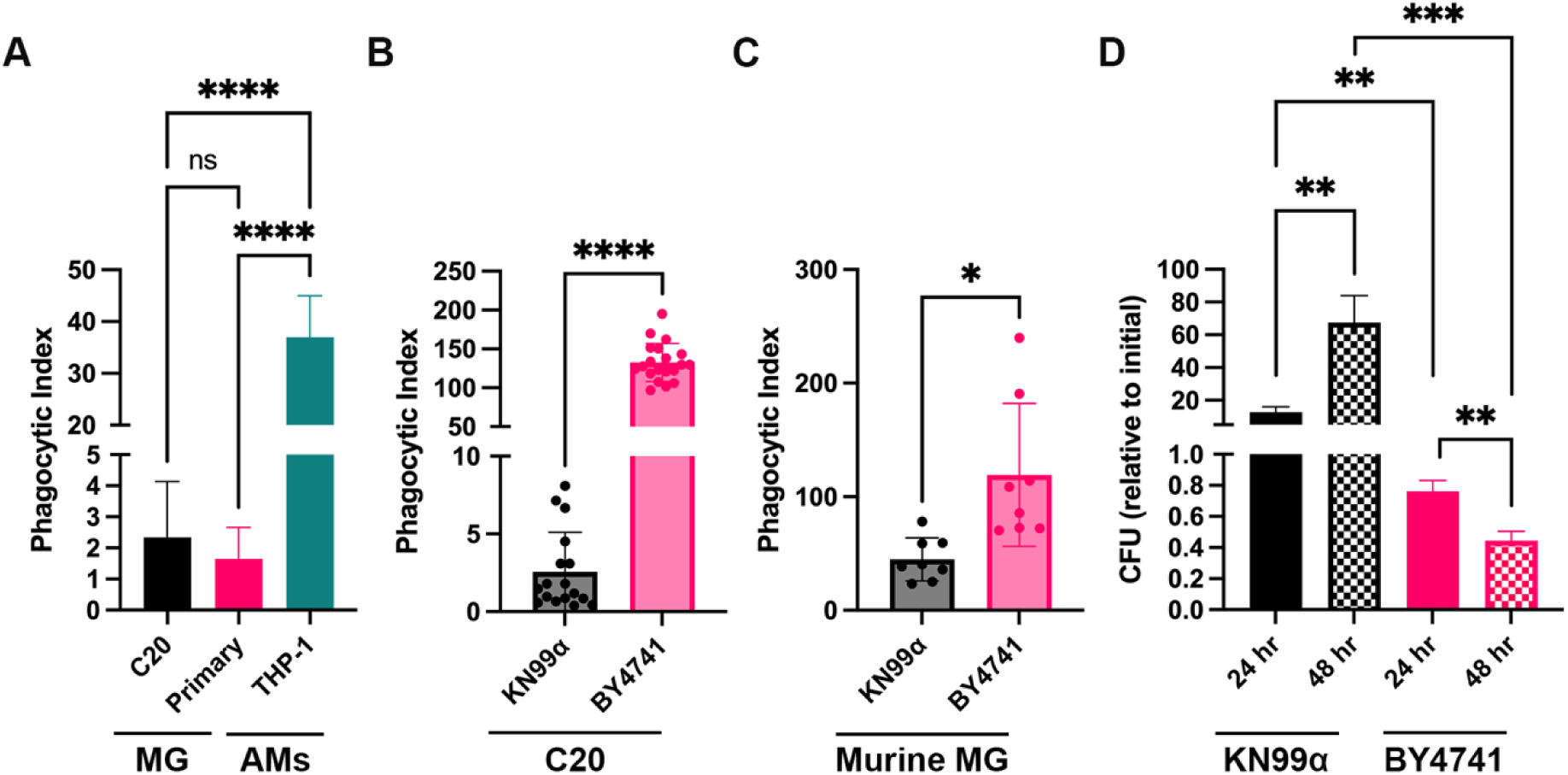
Human microglia are defective at phagocytosing *C. neoformans* and permissive to its intracellularly replication. (A - C) Uptake assay between different macrophages to determine phagocytic indices of WT *C. neoformans* (KN99α) and non-pathogenic *S. cerevisiae* (BY4741). (A) Two types of human microglia (MG) were tested against THP-1 cells, a model of alveolar macrophages (AMs). mCherry-expressing KN99α was serum-opsonized and incubated with the different macrophages at an MOI of 20:1 for 3 hr. Imaging was performed using automated microscopy and analyzed using a CellProfiler pipeline. Values represent the mean ± SD from three biological replicates. Significance was determined using one-way ANOVA with multiple comparisons (Brown-Forsythe and Welch’s corrected), ****, P < 0.0001. (B, C) Uptake assays using C20 immortalized human microglia (B) and primary murine microglia (C) to test engulfment of KN99α and BY4741. mCherry-expressing fungi were serum-opsonized and incubated with the different microglia for 3 hr at an MOI of 20:1. Imaging and analysis were performed as above. Values represent the mean ± SD of three (B) or two (C) biological replicates. Significance was determined using Welch’s t test, *, P < 0.05; ****, P < 0.0001. (D) *In-vitro* survival of KN99α and BY4741 in C20 cells. Fungi and C20 cells were coincubated for 3 hr, at which point one-third of the samples were lysed to determine intracellular fungi, and the other two-thirds were grown for 24 hr and 48 hr, after which they were also lysed. Shown are the colony forming units (CFUs) that were obtained at each timepoint, normalized to the CFUs of initial engulfment (3 hr). Values represent the mean ± SEM from ten biological replicates. Significance was determined using one-way ANOVA with multiple comparisons (Brown-Forsythe and Welch’s corrected), **, P < 0.005; ***, P < 0.0005.

### 2.2. *C. neoformans* evasion of phagocytosis by microglia is not dependent on fungal viability, secreted factors, or capsule production

We next sought to identify factors that could be contributing to the ability of *C. neoformans* to evade phagocytosis by microglia. To do this, we tested a variety of factors that could influence microglial recognition and engulfment of KN99α. First, we tested the ability of C20 cells to engulf live versus heat-killed KN99α (Fig. 2A). We found no significant difference in the PI between viable (alive) or non-viable *C. neoformans* (heat-killed), with heat-killed KN99α only having an average PI fold-change of 2.2 compared to live fungi (normalized to a PI of 1). Next, we looked at the ability of secreted factors, present in conditioned media (CM), to inhibit recognition and phagocytosis by C20 cells. To do this, we prepared KN99α and BY4741 CM and added the BY4741-CM to live KN99α, and KN99α-CM to live BY4741 (in place of basal media, BM, 50% DMEM:Hams F-12) during co-incubation with C20 cells. If there were secreted cryptococcal metabolites, proteins, or capsular material that could inhibit phagocytosis by microglia, we would expect to see a decrease in phagocytosis of BY4741 when incubated with KN99α CM. However, we saw that adding CM to either KN99α or BY4741 did not alter their PI (Fig. 2B). Specifically, when adding BY4741-CM to KN99α, the PI decreased from 1 to 0.43, with this decrease not being statistically significant. Similarly, adding KN99α-CM to BY4741 increased the PI slightly from 38 to 42, with this increase having no statistical significance. Finally, we looked at the impact of capsule production/structure on cryptococcal engulfment by microglia. The polysaccharide capsule of *C. neoformans* is the yeast’s main virulence factor and has been extensively described as antiphagocytic, especially in regard to cryptococcal-macrophage interactions (29–31). Furthermore, the capsule, and its main component (GXM), has been implicated in contributing to the pathogenesis of cryptococcal meningitis (32–34). Thus, we used various mutants that are all defective in capsule organization, structure, or production, and determined the ability of these strains to be phagocytosed by C20 cells. Surprisingly, we found that these mutants have no significant difference in phagocytosis by C20 cells compared to WT (Fig. 2C). Notably, *cap59*Δ, which lacks capsule production altogether, only had a PI 1.4 times that of KN99α. Overall, this data suggests that components other than secreted factors, capsule, or viability are responsible for *C. neoformans*’ ability to evade engulfment by microglia.

**Fig. 2.**
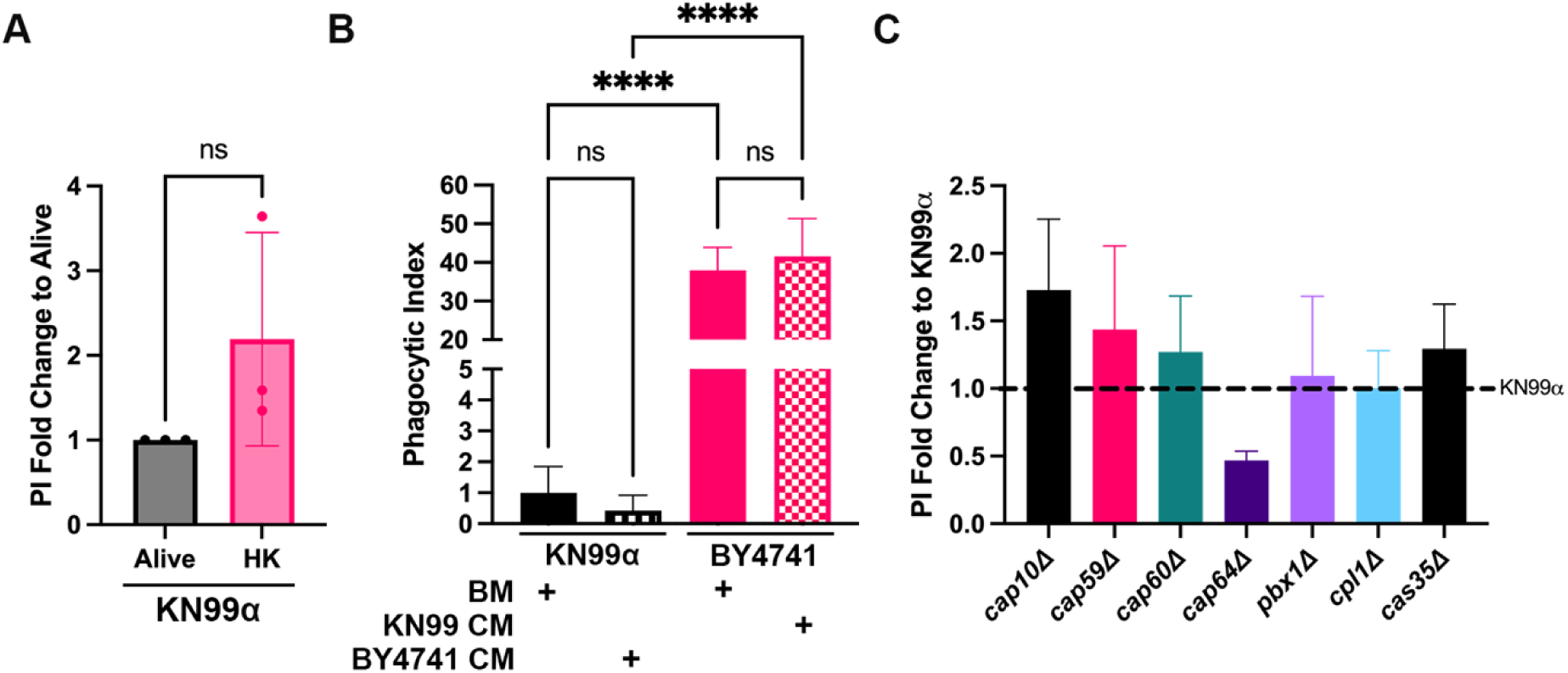
The ability of *C. neoformans* to be recognized and engulfed by C20 cells is independent of fungal viability, secreted factors, or capsule production/structure. (A) An uptake assay showing the ability of C20 cells to engulf live or heat-killed (HK) WT *C. neoformans* (KN99α). Fungi were heat-killed at 75 °C for 30 min. The Y axis denotes the phagocytic index (PI) fold-change to Alive KN99α. Values represent the mean ± SEM from three biological replicates. Significance was determined using Welch’s t test, ns = not significant. (B) Uptake assay using conditioned media (CM) from fungal strains. CM was obtained by growing fungi for 72 hr in 50% DMEM:Hams F-12, followed by centrifugation and filtration of the supernatant. Fungi were coincubated with C20 cells in either basal media (BM; 50% DMEM:Hams F-12) or the opposite fungi’s CM (i.e., KN99α incubated in BY4741-CM, and vice versa). Values represent the mean ± SEM from three biological replicates. Significance was determined using one-way ANOVA with multiple comparisons (Brown-Forsythe and Welch’s corrected), ****, P < 0.0001. (C) Uptake assay comparing the phagocytic index of capsular mutants to KN99α in C20 cells. Mutants were obtained from the Santiago-Tirado lab personal deletion library and the Madhani deletion collection. Y axis denotes the PI fold-change to WT. Significance was determined using one-way ANOVA with multiple comparisons (Brown-Forsythe and Welch’s corrected); none of the strains tested were significant.

### 2.3. Microglia recognize *C. neoformans* differently than peripheral macrophages

To identify cryptococcal proteins that could be contributing to the evasion of phagocytosis by microglia, we assessed the ability of C20 cells to engulf *C. neoformans* mutants that have shown increased engulfment by or altered survival within peripheral macrophages (35–42). As expected, two of these mutants, *pbx1*Δ and *cdk8*Δ show increased internalization by AMs compared to WT (PI = 35.1), with PIs of 53.9 and 52, respectively; however, these mutants show no difference in their phagocytosis levels by C20 cells (PIs of 1.7, 1.2, 1.6 for WT, *pbx1*Δ and *cdk8*Δ, respectively; Fig. 3A). Furthermore, a variety of mutants reported to be high uptake (such as *pdr6*Δ, *ctr2*Δ, *nrg1*Δ, *rim20*Δ, *rdi1*Δ, *blp1*Δ, and *app1*Δ) also showed no differences in their ability to be recognized and engulfed by C20 cells (Fig. 3B). To do a more extensive screen to identify antiphagocytic proteins relevant to microglia-cryptococcal interactions, we tested 50 of 56 mutants identified by Santiago-Tirado *et al*. (41) with altered internalization by AMs (we were unable to recover the remaining 6 from the frozen stocks). Almost all of the 50 mutants exhibited a different phenotype in C20 relative to THP-1 cells (Fig. 3C-D). These results were also compared to those reported by Gaylord *et al*. in bone marrow-derived macrophages (BMDMs) (37). In Santiago-Tirado *et al*., mutants with significantly altered internalization by THP-1 compared to WT were determined by having a PI two standard deviations away from the WT mean. In Gaylord *et. al*., they determined hits with altered internalization by BMDMs by comparing uptake scores (PI fold-change to WT) to the median uptake score for each plate of the deletion collection, then used permissive cut offs to identify 10% of the strains tested as hits. For the C20 screen described in this paper, we tested 9 mutants alongside WT at a time for 1 biological replicate, then repeated this for at least 3 biological replicates for all 50 mutants. The average PI values of each replicate (including mutants and WT) were compiled and One-way ANOVA with multiple comparisons to WT was performed to assess significance between the 50 mutants. Of the 22 mutants tested with increased internalization by THP-1 (green; high uptake (HU)), 12 of these also show increased internalization by BMDMs (Fig. 3C). In contrast, only 3 mutants that show increased internalization by THP-1 shared this antiphagocytic property with C20s (*rim101*Δ, *hpi1*Δ, *hir1*Δ; Fig. 3C). Of the 28 mutants tested with decreased internalization by THP-1 (red; low uptake (LU)), none resulted in decreased internalization by BMDMs or C20 cells (Fig. 3D). Surprisingly, 3 of these showed the opposite phenotype: *trs130*Δ and *lpi1*Δ were high uptake in BMDMs, and *opt1*Δ was high uptake in C20 cells, relative to WT (Fig. 3D). Overall, there was significantly more similarities between THP-1 and BMDMs than between either of them and C20s. Taken together, this indicates that microglia recognize *C. neoformans* differently than peripheral macrophages, and this difference is exacerbated in mutants with increased internalization by peripheral macrophages.

**Fig. 3.**
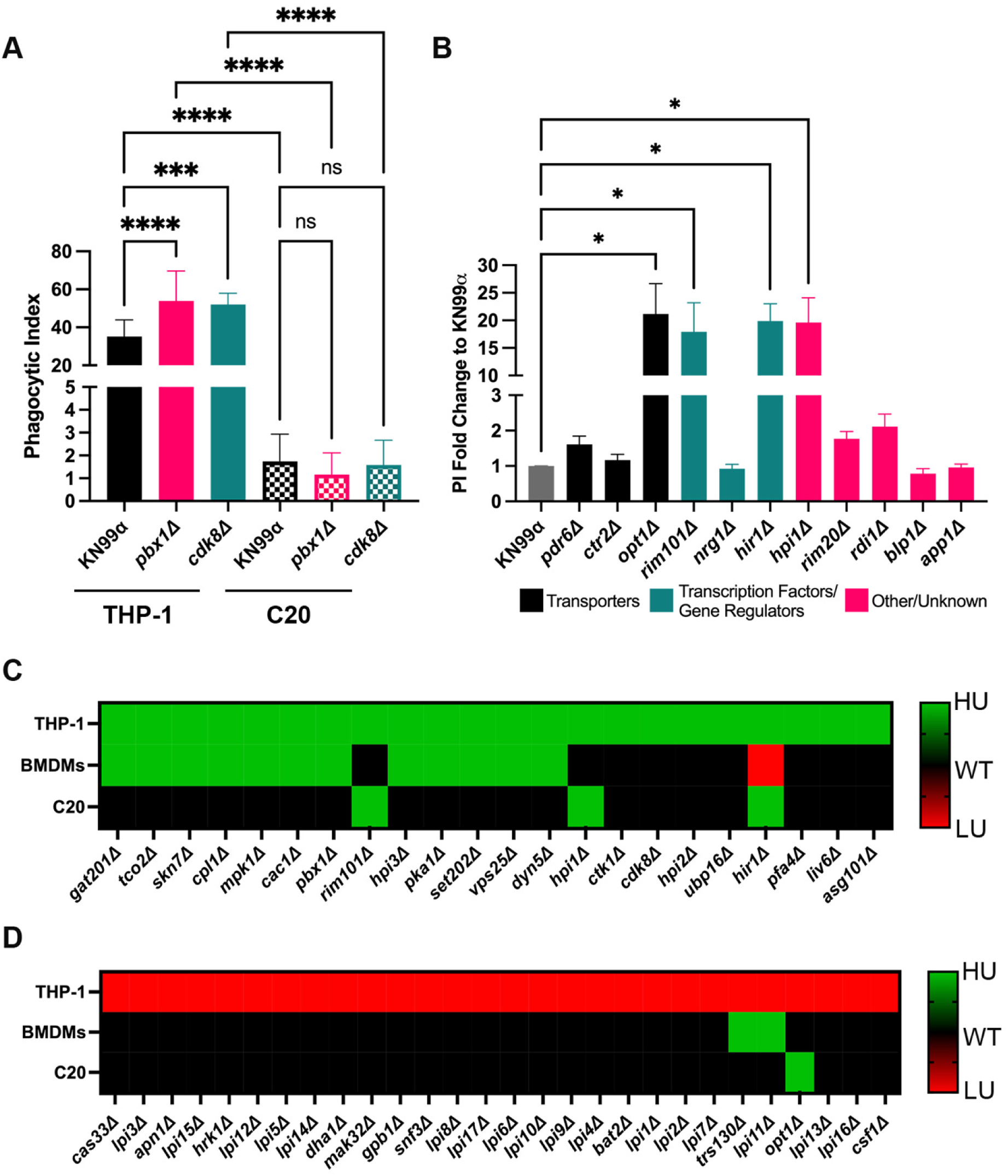
*C. neoformans* recognition and engulfment is distinct between microglia and other macrophages. (A) Uptake assay showing the ability of AMs (THPs) and immortalized human microglia (C20) to engulf WT and mutant strains of *C. neoformans*. The mutants tested were characterized by Santiago-Tirado et al. as having increased internalization by AMs and were obtained from the Madhani deletion collection. Values represent the mean ± SEM from three biological replicates. Significance was determined using one-way ANOVA with multiple comparisons (Brown-Forsythe and Welch’s corrected), ***, P < 0.0005; ****, P < 0.0001. (B) Uptake assay showing the PI as fold-change to WT of different cryptococcal mutants when coincubated with C20 cells. The mutants tested were described in previous literature as having increased internalization or altered interactions with macrophages. The mutants were obtained from the Madhani deletion collection. Values represent the mean ± SEM from three biological replicates. Significance was determined using one-way ANOVA with multiple comparisons (Brown-Forsythe and Welch’s corrected), *, P < 0.05. (C, D) Results from uptake assays done with C20 cells (HMG) and various cryptococcal mutants and compared to the results of screens done in THPs (AMs) and bone marrow-derived macrophages (BMDMs) that were identified in the literature. The mutants tested were screened by Santiago-Tirado et al. and identified as those with increased (C) and decreased (D) internalization by AMs. Green represents mutants with increased internalization by the specified macrophage (as determined from the thresholds set forth by the various authors of the publications chosen). Red represents mutants with decreased internalization, and black represents mutants with no significant altered interactions with macrophages when compared to WT. For the C20 screen, results indicate values that represented the mean ± SEM from three biological replicates (the values for the mutants with enhanced internalization by HMG are shown in panel B). Significance was determined using one-way ANOVA with multiple comparisons (Brown-Forsythe and Welch’s corrected), *, P < 0.05.

### 2.4. *C. neoformans* disrupts phagosome maturation in human microglia

It has been shown in various types of peripheral macrophages that *C. neoformans* manipulates the phagosome maturation process by delaying association of maturation markers (16, 21, 22). Since we show that microglia have altered interactions with *C. neoformans* compared to AMs, we wanted to study the phagosome maturation process of KN99α in human microglia after engulfment (Fig. 4). To do this, we performed immunofluorescence using primary antibodies against phagosome maturation markers (EEA1, LAMP1, vATPase) to assess percent-positive cargo-containing phagosomes after 1, 2, and 3 hours of incubation with either C20 cells or primary human microglia. EEA1 is a marker for early phagosomes, and we found that our controls, BY4741 and 5 μm latex beads, had significantly more phagosomes with EEA1 association at earlier timepoints in both C20 cells and primary human microglia than KN99α (Fig. 4A-B). Furthermore, EEA1 association follows the same trend for BY4741 and latex beads, with the number of percent-positive EEA1-associated phagosomes decreasing over time in C20 cells and primary human microglia (from 95.1 and 89.7% to 66.8 and 51.6%, respectively, in C20 cells, and from 96.2 and 93.8% to 70.2 and 57.3%, respectively, in primary microglia). However, in C20 cells and primary human microglia, the percentage of EEA1-associated KN99α-containing phagosomes increases during the timecourse (from 64.9 to 76.6% in C20 cells and 65.9 and 87.4% in primary human microglia). This indicates that KN99α not only delays EEA1 association to its phagosome but retains it in both immortalized and primary human microglia. vATPase accumulates on phagosomes as they progress through the maturation pathway and is the primary enzyme responsible for phagosome acidification. In C20 cells and primary human microglia, all cargo-containing phagosomes increase in vATPase association during the 3-hour timecourse; however, KN99α has a significant decrease in vATPase-positive phagosomes at all timepoints compared to BY4741 and latex beads (Fig. 4C- D). Specifically, at 3 hours, in C20 cells and primary human microglia, the percentage of KN99α-containing phagosomes positive for vATPase were 64.7 and 50.8, respectively, compared to BY4741-containing phagosomes, which were 94 and 89.7, respectively. This pattern is also seen with cargo-containing phagosomes positive for LAMP1 association. LAMP1 is a marker for late-stage phagosomes and lysosomes, and routinely used as a marker for phagolysosomes. In both types of microglia, all cargo-containing phagosomes increase in LAMP1 association over time (Fig. 4E, F). However, in both cell types, KN99α-containing phagosomes have significantly lower LAMP1 association than BY4741 and latex beads phagosomes at all timepoints (Fig. 4E, F). Taken together, this data indicates that KN99α-containing phagosomes have delayed EEA1, LAMP1, and vATPase association, indicating altered phagosome maturation in immortalized and primary human microglia. The similar trends for all phagosome maturation markers across microglia cell types is also an indicator of the translatability of the C20 cell line to that of primary cells, especially as it relates to studying cryptococcal infection.

**Fig. 4.**
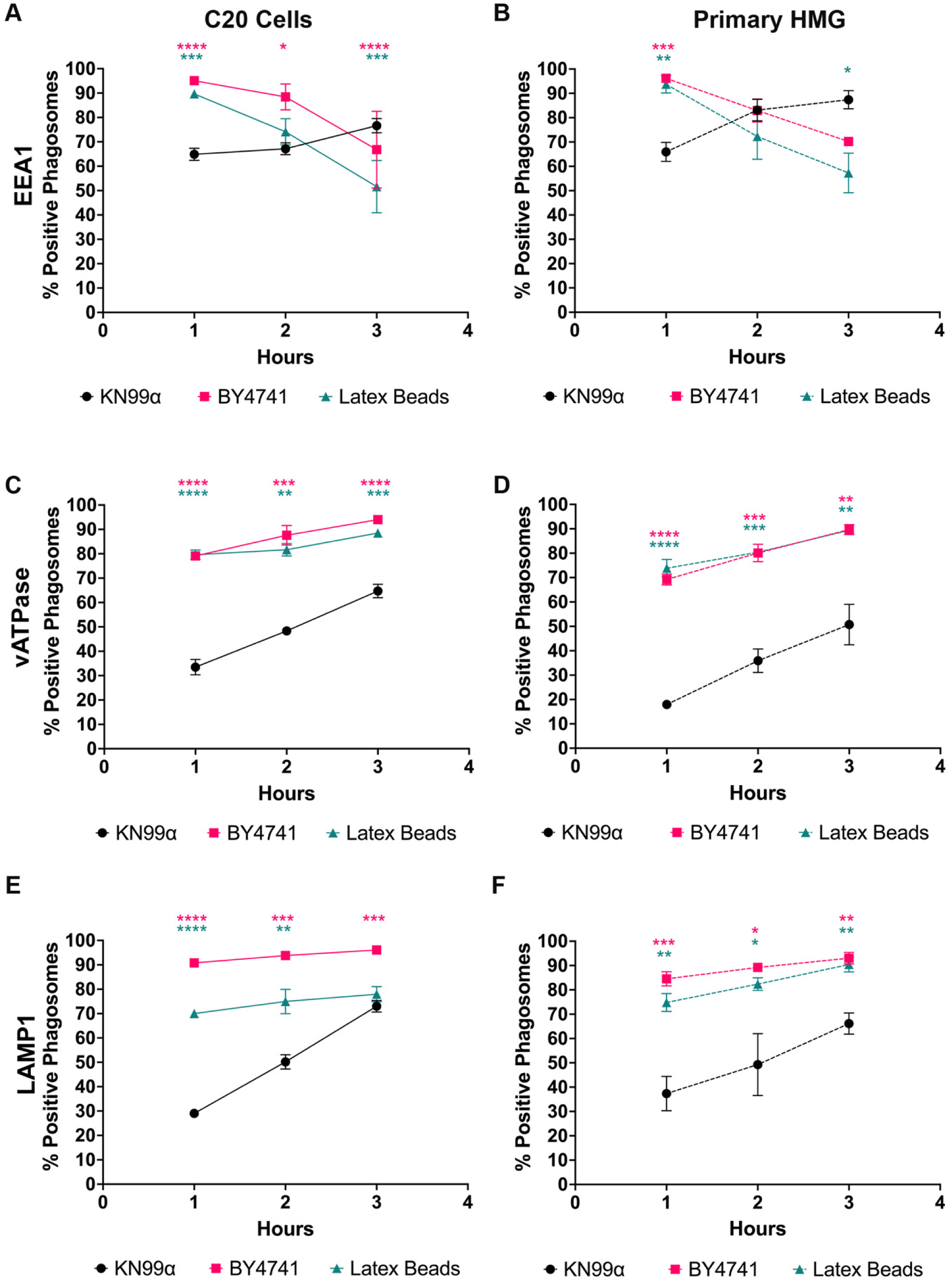
Maturation of cryptococcal phagosomes in immortalized and primary human microglia is altered. (A, B) Immunofluorescence showing the cargo-containing phagosomes that are positive for EEA1 association at 1, 2, and 3 hours post-coculture with C20 cells (A) or primary human microglia (B). mCherry-expressing fungi or fluorescent latex beads were opsonized and incubated with microglia at an MOI of 20:1 (fungi) or 10:1 (latex beads) for 1, 2, or 3 hours. Coverslips were fixed, stained with primary and secondary antibodies, and imaged via fluorescent microscopy. (C, D) Plotted are the cargo-containing phagosomes that are positive for vATPase association at 1, 2, and 3 hours post-coculture with C20 cells (C) or primary human microglia (D). (E, F). Plotted are the cargo-containing phagosomes that are positive for LAMP1 association at 1, 2, and 3 hours post-coculture with C20 cells (E) or primary human microglia (F). Intracellular cargo shown are WT *C. neoformans* (KN99α), and *S. cerevisiae* (BY4741) and 5 um red fluorescent latex beads (CD Bioparticles) as controls. Lines representing C20 cells are solid, and dashed lines represent primary human microglia. Values represent the mean ± SEM from three biological replicates. Significance was determined using one-way ANOVA with multiple comparisons (Brown-Forsythe and Welch’s corrected) for each timepoint, *, P < 0.05; **, P < 0.005; ***, P < 0.0005; **** P < 0.0001.

### 2.5. Different cryptococcal strains alter phagosome maturation in microglia

Since we show that KN99α alters phagosome maturation in both immortalized and primary human microglia, we wanted to test the same with mutants with known altered *in-vitro* and/or *in-vivo* survival in peripheral macrophages (Fig. 5). *Cap59*Δ lacks capsule production and has been shown to be avirulent in various animal models (43–45). At all timepoints, *cap59*Δ has a significantly higher proportion of LAMP1-positive phagosomes compared to KN99α, indicating that this mutant has normal progression through the phagosome maturation pathway (Fig. 5A). Additionally, *cap59*Δ has significantly lower CFUs after 24- and 48-hours post-engulfment compared to KN99α (Fig. 5A). While the normalized CFU counts for *cap59*Δ are >1, indicating intracellular replication, this data supports that C20 cells have enhanced fungistatic activity against this mutant. *Pdr6*Δ is a mutant that has been shown to have altered survival within macrophages and a mouse model (42, 46). In C20 cells, this mutant has increased LAMP1 association with phagosomes (albeit still at a lower rate than controls) at 1- and 2-hours compared to KN99α; however, this accelerated LAMP1 association did not lead to a significant reduction in normalized CFU count after 24- or 48-hours post engulfment (Fig. 5B). *Rdi1*Δ has difficulty surviving and proliferating within macrophage phagosomes and has attenuated virulence in a mouse model (40). We find that *rdi1*Δ has similar LAMP1-association and intracellular survival compared to *pdr6*Δ, with significantly higher LAMP1-positive phagosomes at 1- and 2-hours compared to WT, and no significant reduction in normalized CFU count after 24- or 48-hours post-engulfment (Fig. 5C). In a mouse infection model, *gat201*Δ is undetectable in lungs by 10-dpi, indicating fungal clearance (35). We find that *gat201*Δ has significantly increased LAMP1-positive phagosomes at 1- and 2-hours, and significantly reduced internalized CFU counts at 48-hours (Fig. 5D). *Mpk1*Δ has attenuated virulence in a mouse model (47), and similar to the other mutants tested, *mpk1*Δ has accelerated LAMP1 association to its phagosomes compared to WT at 1- and 2-hours, and reduced internalized CFU after 48-hours, indicating enhanced fungistatic activity of C20 cells (Fig. 5E). Lastly, *VPS25* encodes a member of the ESCRT complex, and its mutant (as well as mutants missing other proteins within this complex) has been shown to have altered survival within macrophages and attenuated virulence in mouse models (48). Following a similar trend to *gat201*Δ and *mpk1*Δ, *vps25*Δ has accelerated LAMP1 association to its phagosomes at 1- and 2- hours and has significantly decreased normalized CFUs at 48-hours post-engulfment by C20 cells (Fig. 5F). This indicates that C20 cells have enhanced fungistatic activity against *gat201*Δ*, mpk1*Δ, and *vps25*Δ. Taken together, all of the mutants tested have accelerated LAMP1 association to their phagosomes shortly after engulfment by C20 cells, however, this does not correlate with microglia fungicidal activity.

**Fig. 5.**
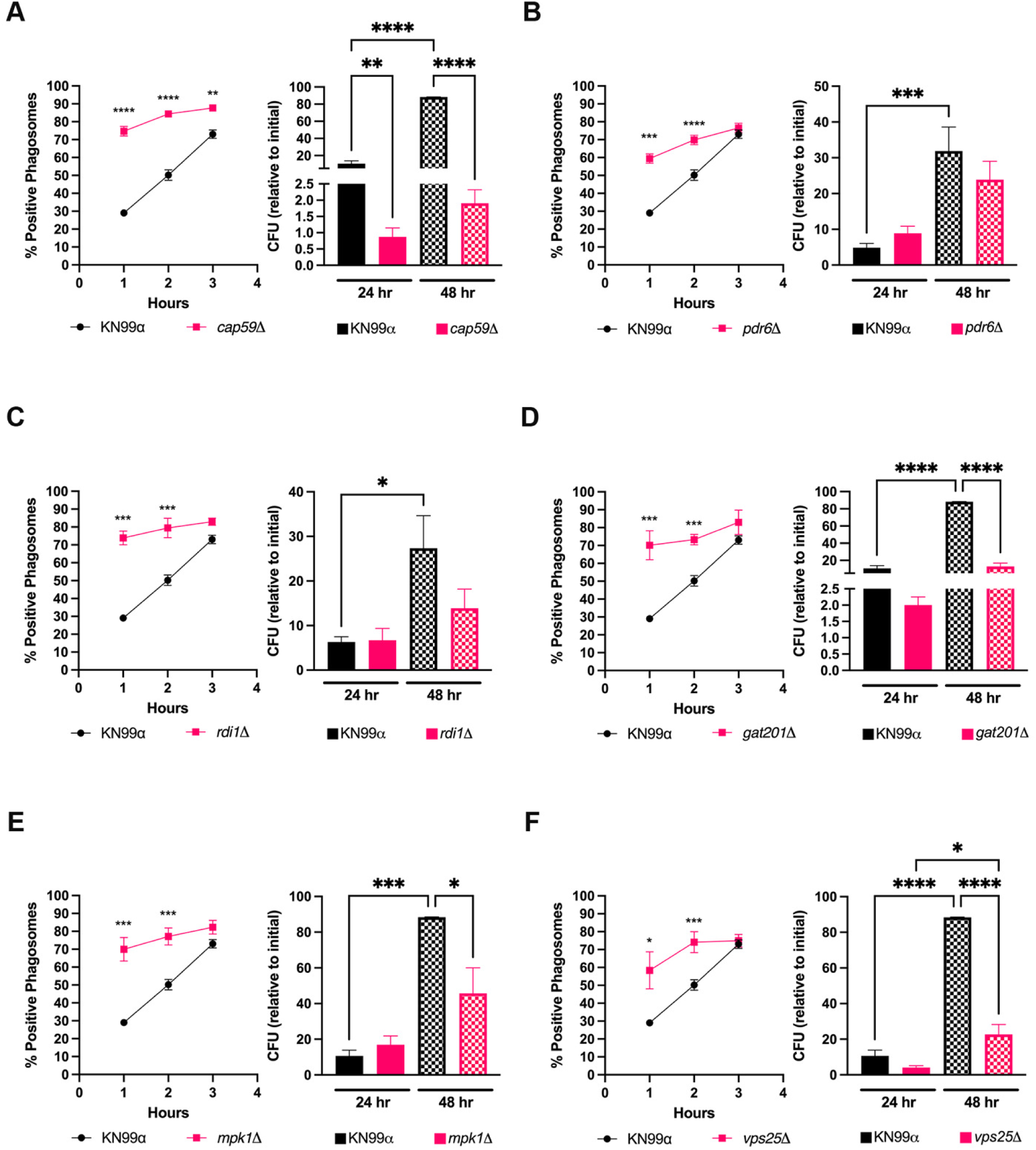
Phagosome maturation alterations in microglia does not completely correlate with fungicidal activity. (A) Immunofluorescence showing the KN99α- and *cap59*Δ-containing phagosomes that are positive for LAMP1 association at 1, 2, and 3 hours post-coculture with C20 cells (left). *In-vitro* survival assay assessing internalized KN99α and *cap59*Δ after 24- and 48-hr post-engulfment by C20 cells (right). Shown are the colony forming units (CFUs) that were obtained at each timepoint, normalized to the CFUs of initial engulfment (3 hr). (B-F) representation of experiments performed in panel A, however, with KN99α and *pdr6*Δ (B), *rdi1*Δ (C), *gat201*Δ (D), *mpk1*Δ (E), and *vps25*Δ (F). The mutants chosen for this figure all have been characterized as having altered survival within macrophages. Values represent the mean ± SEM from three biological replicates. For all experiments, significance was determined using one-way ANOVA with multiple comparisons (Brown-Forsythe and Welch’s corrected), *, P < 0.05; **, P < 0.005; ***, P < 0.0005; ****, P < 0.0001.

### 2.6. *C. neoformans* induces phagosome membrane damage in human microglia

Galectin-3 (Gal-3) staining is widely used as a reporter of lysosomal membrane damage because it will bind the sugars on the luminal side of phagosomes only if membrane integrity is lost (49, 50). Using Gal-3 staining, we have shown that *C. neoformans* induces phagosome membrane damage in AMs as early as 4 hours post-infection (21). Thus, we wanted to determine if *C. neoformans* also induces phagosome membrane damage in microglia. To do this, we incubated *C. neoformans* or *S. cerevisiae* with C20 cells for 3 hours to allow for fungal engulfment. Extracellular fungi were washed, and the infected microglia were incubated for an additional 24 hours, and their phagosomes were assessed for membrane damage via immunofluorescence using an antibody against Gal-3. We found that both unopsonized and serum-opsonized KN99α and BY4741 had phagosomes positive for Gal-3 after 24 hours (Fig. 6A). However, KN99α had significantly more Gal-3-positive phagosomes compared to BY4741, with an average of 41.3% and 15.6%, respectively, for unopsonized fungi. For serum-opsonized fungi, KN99α had 58.5% phagosomes positive for Gal-3 compared to 26.9% for BY4741. Furthermore, serum-opsonized KN99α-containing phagosomes had significantly more Gal-3 association than phagosomes containing unopsonized KN99α (58.5% and 41.3%, respectively). Overall, this data indicates that microglia are extremely sensitive to phagosome permeabilization by *C. neoformans* at 24 hours after engulfment.

**Fig. 6.**
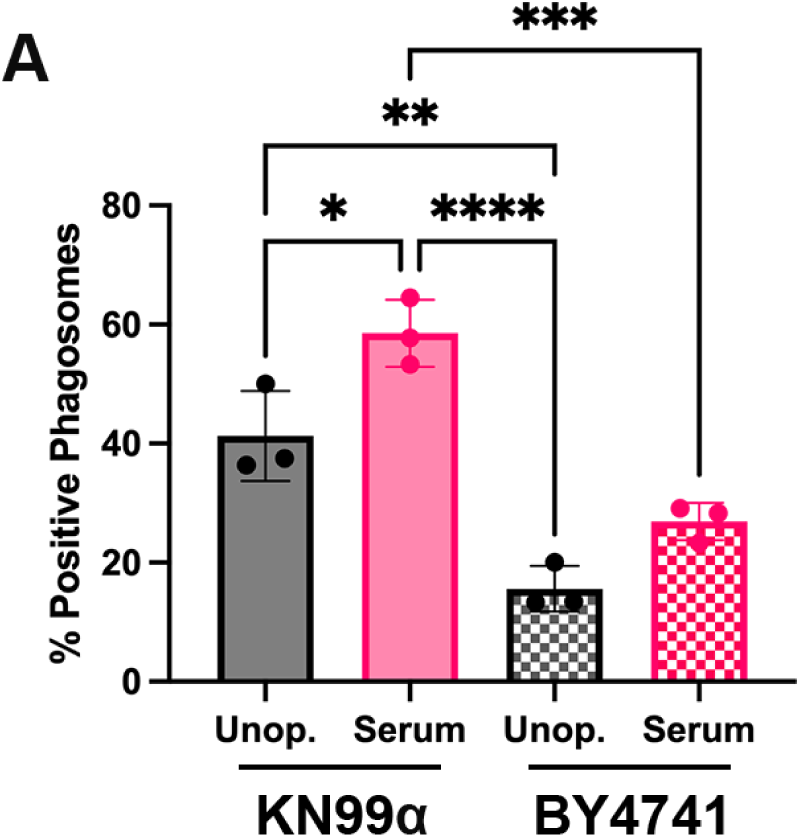
Phagosome membrane damage occurs in cryptococcal-containing phagosomes. (A) Immunofluorescence showing the percentage of fungal-containing phagosomes positive for Gal-3 association 24 hr after engulfment. Gal-3 is a marker used for phagosomal membrane damage. Both *C. neoformans* (KN99α)- and *S. cerevisiae* (BY4741)-containing phagosomes were tested for membrane damage. mCherry-expressing fungi were either unopsonized (unop.) or opsonized (serum) and incubated with microglia at an MOI of 20:1 for 3 hr. Wells were washed three times with DPBS to remove extracellular fungi, and microglia were incubated for an additional 24 hr then assessed for membrane damage via Gal-3 staining. Values represent the mean ± SEM from three biological replicates. For all experiments, significance was determined using one-way ANOVA with multiple comparisons (Brown-Forsythe and Welch’s corrected), *, P < 0.05; **, P < 0.005; ***, P < 0.0005; ****, P < 0.0001.

### 2.7. Cryptococcal strains with increased internalization by microglia exhibit altered cell body size and cell wall composition

Thus far we have shown that *C. neoformans* evades phagocytosis by microglia and, if internalized, can manipulate phagosome maturation, and that these processes can be altered by various mutants. Therefore, we wanted to study mutants that have increased engulfment by microglia to identify factors that could be attributed to their altered recognition. We chose to study the 4 mutants identified as high-uptake in Fig. 3 for variations in capsule production, cell body size, and cell wall structure (Fig. 7). We first assessed the ability of these mutants to produce capsule (Fig. 7A-B). The only mutant that had altered capsule production compared to WT was *hpi1*Δ, which on average had a significantly larger capsule radius (Fig. 7B). We then assessed the cell body size of these mutants in host-like conditions (DMEM, 37°C, 5% CO_2_). All mutants except for *hir1*Δ had significantly smaller cell bodies compared to WT (Fig. 7C). Similarly, when assessing the cell body size of these mutants grown in nutrient-rich medium (YPD), all 4 mutants had significantly smaller cell bodies (Fig. 7D). Overall, this data indicates that cryptococcal mutants with increased internalization by microglia generally have a smaller cell body size compared to WT. Lastly, we assessed the cell wall composition of the 4 mutants by staining with dyes known to bind specific components of the cell wall (Fig. 7E-G). All mutant strains had significantly increased staining of calcofluor white (CFW) and eosin Y (EoY), which are indicative of enhanced chitin and chitosan exposure, respectively. We also stained with concanavalin A (ConA) to assess mannoprotein exposure, and found that only one mutant, *rim101*Δ, had significantly increased ConA staining compared to WT. Overall, this data indicates that enhanced chitin and chitosan exposure could contribute to enhanced recognition and engulfment by C20 cells.

**Fig. 7.**
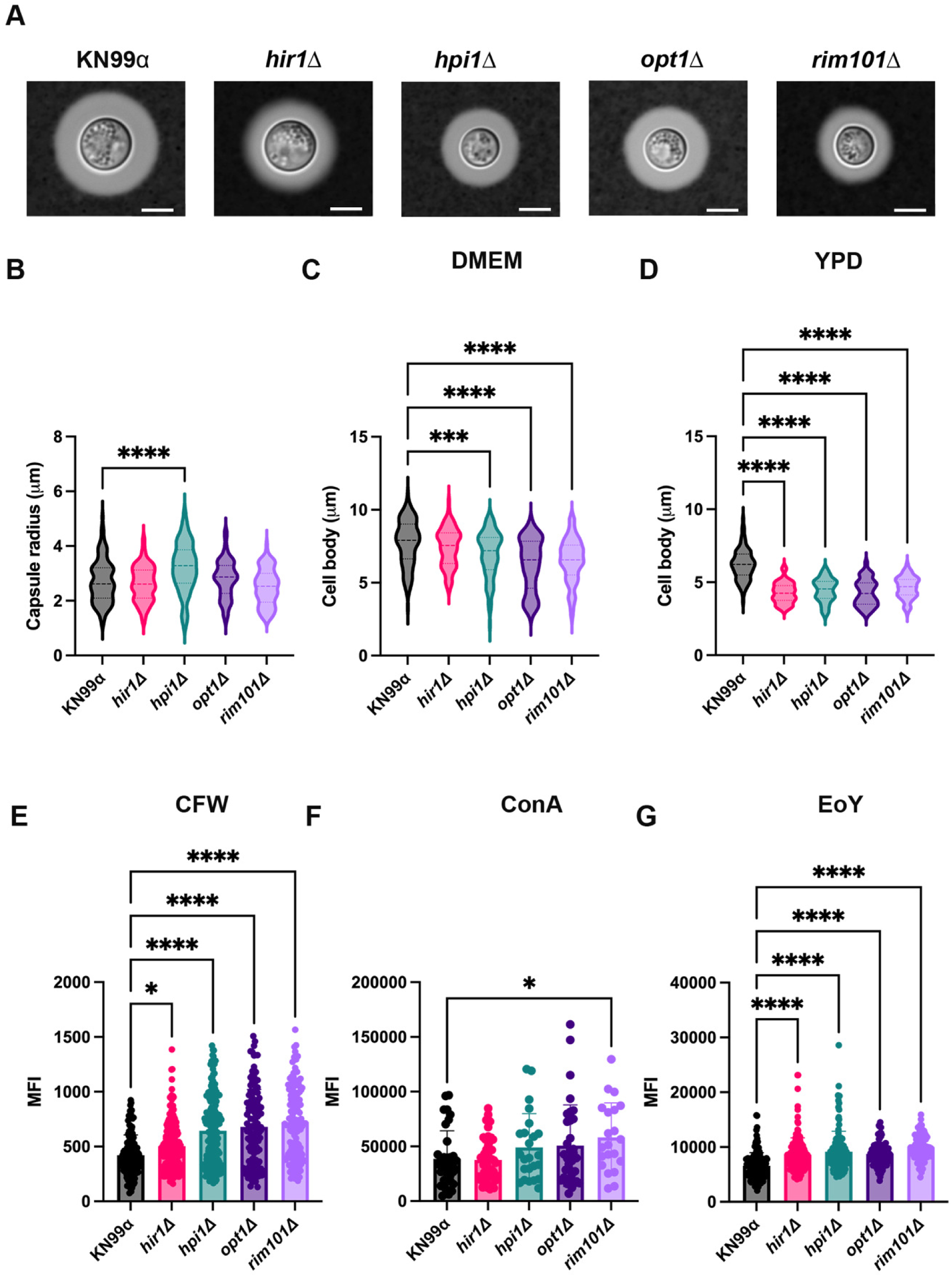
Cryptococcal cell body size and cell wall composition could attribute to recognition and engulfment by microglia. (A) Representative capsule induction images of WT *C. neoformans* (KN99α) and the four mutants identified as having increased internalization by C20 cells in Fig. 3 (*hir1*Δ, *hpi1*Δ, *opt1*Δ, and *rim101*Δ). Capsules were induced by growing the fungal strains in DMEM for 24 hr and imaged at 100X after India Ink staining. The light gray shading around the cell body is the polysaccharide capsule. Scale bar represents 5 μm. (B) Quantification of the capsule radius in μm from the images represented in panel A. Capsule radius analysis determined via ImageJ software. Equation for capsule radius is (total cell size including capsule - cell body size)/2. (C) Quantification of the cell body size in μm from the images represented in panel A. Cells were grown in DMEM for 24 hr. Measurements taken using ImageJ software (D) Quantification of the cell body size in μm from cells grown in YPD for 18 hr. Measurements taken using ImageJ software. (B-D) Values are represented as a violin plot showing all measurements (< 100 cells) from three biological replicates. (E-G) Quantification of mean fluorescent intensity (MFI) of fungal cells stained with calcofluor white (CFW; E), concanavalin A (ConA; F), or eosin Y (EoY; G). Fungal cultures were washed with PBS and stained with their respective cell wall dyes for 30 min. Cells were washed three times with PBS and imaged via fluorescent microscopy. Quantification of MFI was determined via Cell Profiler pipeline. Values represent the mean ± SD from three biological replicates. For all experiments, significance was determined using one-way ANOVA with multiple comparisons (Brown-Forsythe and Welch’s corrected), *, P < 0.05; ***, P < 0.0005; ****, P < 0.0001.

### 2.8 Assessment of additional virulence factors in mutants with enhanced engulfment by microglia

Since we already tested the ability of *hir1*Δ, *hpi1*Δ, *opt1*Δ, and *rim101*Δ to produce capsule, we wanted to perform experiments assessing additional virulence factors of *C. neoformans*. Specifically, we tested melanization, urease production, growth at high temperature, and resistance to oxidative stress (Fig. 8). First, we incubated the cryptococcal strains on melanin-inducing solid media and found that *hir1*Δ qualitatively produces more melanin than WT (Fig. 8A). All other mutants had similar melanin production to WT. *Lac1*Δ was used as a negative control, as this mutant lacks the gene responsible for melanin production. We also tested the ability of these strains to retain melanin in their cell wall. To do this we grew the strains in melanin-inducing liquid media overnight and pelleted the cultures. We measured the OD_475_ of the supernatant to quantitate shed melanin. Blanks (non-inoculated liquid L-DOPA media) and *lac1*Δ were used as controls for negative melanin production/shedding. *Hir1*Δ is the only mutant with significantly decreased shed melanin compared to WT, as a higher OD reading is correlated with higher melanin shed into the culture. Overall, this data indicates that *hir1*Δ has cell well alterations that contribute to less shed melanin. This finding may explain the qualitative ‘hypermelanization’ phenotype seen with *hir1*Δ on solid medium, as with less shed melanin, more melanin will accumulate on the cell wall, leading to a darker appearance on solid medium.

**Fig. 8.**
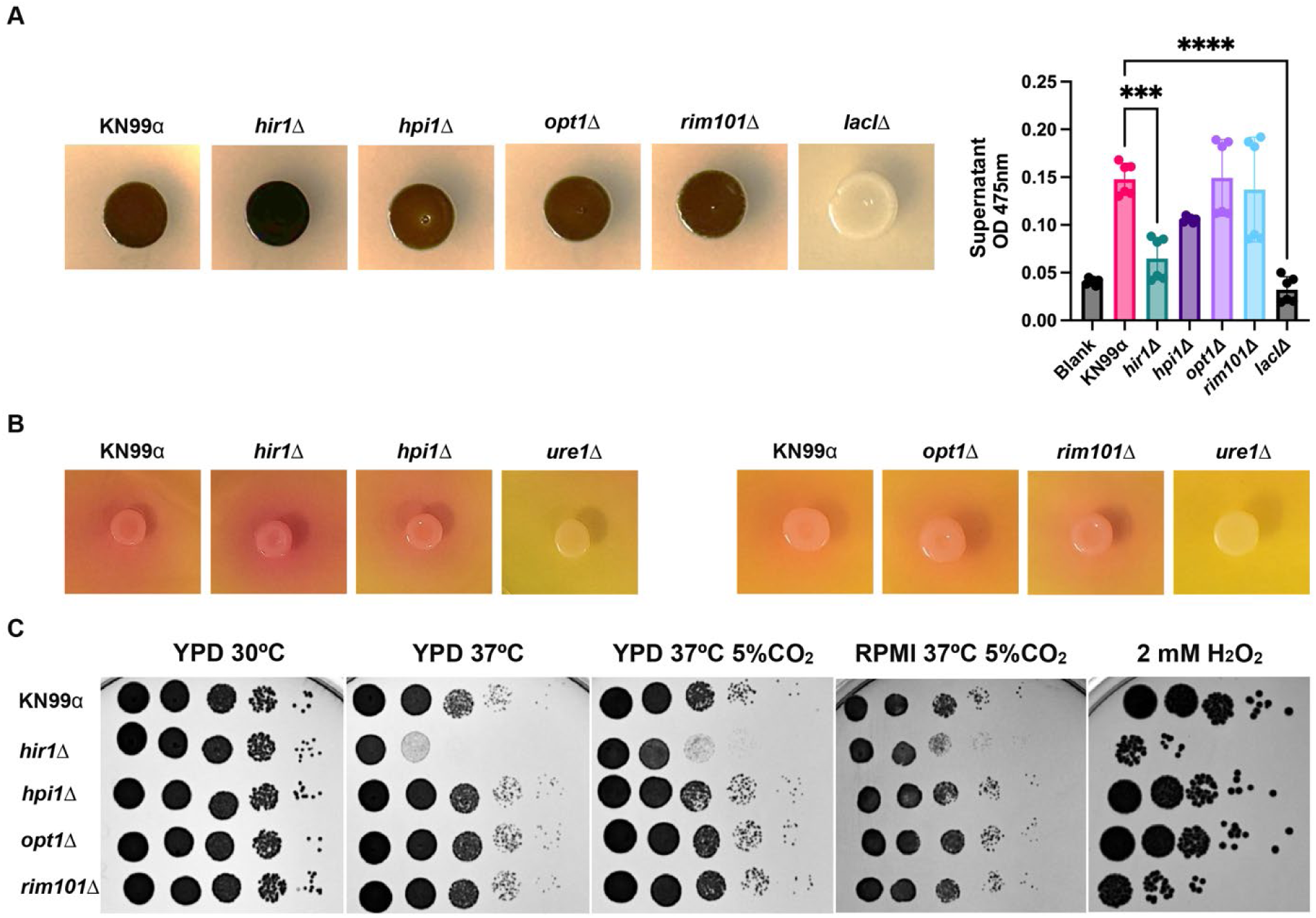
Assessment of stress phenotypes in mutants with high uptake by microglia. (A; left) Representative images of melanin production of WT *C. neoformans* (*neoformans* (KN99α) and the four mutants identified as having increased internalization by C20 cells in Fig. 3 (*hir1*Δ, *hpi1*Δ, *opt1*Δ, and *rim101*Δ). Fungal cultures normalized to an OD_600_ of 0.25 were spotted onto L-DOPA solid media and incubated for 72 hr. (A; right) Measurement of OD_475_ of supernatant of fungal cultures grown in melanin-inducing liquid media. L-DOPA fungal cultures normalized to an OD_600_ of 0.25 were grown for 18 hr, pelleted, and the OD of the supernatant was measured. The liquid L-DOPA media induces melanization, and pelleting the 18 hr cultures allows for visualization of melanin that has shed from the cryptococcal cell wall. A lower OD_475_ indicates less shed melanin. Blank was non-inoculated L-DOPA liquid media included as a control. *LacI*Δ is used as a negative control. Values represent the mean ± SD from three biological replicates. Significance was determined using one-way ANOVA with multiple comparisons (Brown-Forsythe and Welch’s corrected), ***, P < 0.0005; ****, P < 0.0001. (B) Representative images of urease production of the cryptococcal strains. *Ure1*Δ is used as a negative control. Fungal cultures were normalized to an OD_600_ of 0.25 and spotted on Christensen Urea Agar media and incubated for 72 hr. Yellow background indicates no urease production; As urease is produced, the background changes to a pink/red. (C) Growth of cryptococcal mutants assessed under various conditions. Serial dilutions of fungal cultures were prepared and plated on nutrient-rich media (yeast-peptone-dextrose; YPD) and incubated at 30°C, 37°C, or 37°C 5% CO_2_; plated on RPMI agar and incubated at 37°C 5% CO_2_; plated on YPD supplemented with 2 mM H2O2 and incubated at 30°C. All plates were incubated for 48 hr.

The next virulence factor we looked at was urease production. To do this, we plated the strains on urea agar and incubated them for 5 days, including *ure1*Δ as a negative control for urease production (Fig. 8B). As the strains produce urease, the urea in the agar is degraded into ammonia, which increases the pH of the medium leading to a visible color change. Overall, none of the mutants show altered urease production compared to WT.

Lastly, we tested the ability of the *C. neoformans* strains to grow under various stress conditions using agar plates (Fig. 8C). Under optimal growth conditions, such as YPD at 30°C, none of the mutants show a growth defect. However, on YPD at 37°C, *hir1*Δ exhibits a growth defect compared to WT. This defect is partially rescued with the addition of 5% CO_2_. Similarly, *hir1*Δ has a minor growth defect on RPMI media, which mimics host-like conditions. We also tested the ability of the strains to grow under oxidative stress, as this is the main stress the fungus encounters inside a host. When growing these strains on agar plates containing 2 mM H_2_O_2_, we found that *hir1*Δ and *rim101*Δ have increased sensitivity to oxidative stress compared to WT. Overall, this data indicates that *hir1*Δ exhibits a growth defect at high temperature, and two of the four mutants have increased sensitivity to oxidative stress.

## 3. Discussion

Studies of cryptococcal-microglia interactions have lagged behind those focusing on alveolar or peripheral macrophages. Most of what we know regarding microglia and *C. neoformans* comes from *in-vivo* observations in various animal models of *C. neoformans* pathogenesis (14, 51, 52). However, in-depth *in-vitro* interactions studying recognition, phagocytosis, and intracellular survival of *C. neoformans* by human microglia remains widely understudied. Here, we present an initial characterization of cryptococcal-microglia interactions, focusing on phagocytosis evasion, phagosome maturation, and recognition of cryptococcal mutants. We do this using C20 cells, which are a relatively recent, and highly translational, immortalized human microglia cell line (25). We not only define initial cryptococcal-microglia interactions, but also validate the use of C20 cells, when compared to primary human and murine microglia, for studying these interactions *in-vitro*.

We show that immortalized and primary human microglia have similar levels of phagocytosis of WT *C. neoformans*, and we directly compare this with the phagocytic activity of THP-1 cells, a widely used model for AMs. While it has been implied that both human and murine microglia have little to no phagocytic activity against *C. neoformans* (12, 53–55), we are the first to directly compare these phagocytes and show that microglia are significantly defective in internalization of *C. neoformans* relative to AMs. Furthermore, we find that these low levels of cryptococcal engulfment are not due to an inherent defect in phagocytosis, as all C20 cells and primary human and murine microglia have significantly higher internalization of *S. cerevisiae* compared to *C. neoformans*. We have used a variety of microglia (i.e., primary human and murine) to validate the usage of C20 cells to study these fungal interactions, however, we do recognize that primary murine microglia have higher levels of engulfment of WT *C. neoformans* compared to that of C20 cells and primary human microglia. This is not surprising, as many publications have highlighted the differences between rodent-derived and human microglia, specifically noting their distinctions in RNA expression, immunophenotypes, and nitric oxide activity (27, 56–61). This finding does not diminish the credibility of C20 cells, yet only highlights the translatability of this cell line, specifically when comparing it to primary human microglia. We also show, like others have published, that *C. neoformans* survives and replicates inside immortalized human microglia, further showcasing the validity of using the C20 cell line.

Having shown that C20 cells behave just like primary microglia in terms of cryptococcal interactions, we then tried to identify fungal factors that could be mediating these limited levels of engulfment by microglia. While it has been shown that viability is needed for phagosome maturation manipulation by *C. neoformans* (17, 22), we were surprised to see that cryptococcal viability does not, however, impact recognition and engulfment by C20 cells. Furthermore, we find that secreted cryptococcal factors, such as shed capsular material, also do not contribute to phagocytosis evasion by microglia. Similarly, mutants either lacking capsule altogether (*cap59*Δ) or with altered capsule structure/production (*cap10*Δ, *cap60*Δ, *cap64*Δ, *pbx1*Δ, *cpl1*Δ, and *cas35*Δ) also are not engulfed by C20 cells to the same extent as non-pathogenic controls. GXM, the main component of the *C. neoformans* polysaccharide capsule, has been extensively described as antiphagocytic, and plays a significant role during cryptococcal pathogenesis within the CNS (33, 62–64). However, GXM production and shedding is not the only antiphagocytic mechanism employed by *C. neoformans*, as other mechanisms have been described albeit only in peripheral macrophages (35, 65, 66). Thus, it is conceivable that a novel mechanism is at play in the CNS environment, specifically affecting microglia. Although we acknowledge that more testing is necessary, specifically following conditions that induce capsule production, as 3 hour coincubation with C20 cells in DMEM:Hams F-12 might not be sufficient to produce robust capsule and cause a change in phagocytic activity, we believe that C20 cells represent a powerful way to start addressing this question.

Given that C20 cells engulf *S. cerevisiae* cells robustly, we hypothesize that the microglia antiphagocytic phenotype is mediated by cryptococcal proteins. To begin deciphering this, we tested mutants that have been previously characterized as having altered internalization by peripheral macrophages (THP-1 and BMDMs). We found that most of the mutants tested did not reflect the same phagocytic activity by C20 cells that was seen with peripheral macrophages. Notably, there was some concordance between the two peripheral macrophages cell types, however, it is important to highlight the limitations of these comparisons. First, the screens that were done in other macrophages had different thresholds for identifying mutants with altered internalization (these thresholds for THP-1 and BMDMs compared to C20 cells are stated in the Results section). Second, similar to differences between rodent-derived and human microglia, murine-derived BMDMs and human-derived THP-1 have significant differences in their RNA expression, immunophenotypes, and metabolic activity (67–71). Third, while no studies have been done looking at specific differences between microglia and AMs, many studies have looked at the differences in receptor expression and immune responses between microglia and other macrophages within the CNS, including microglia-like cells, which are BMDMs that colonize the CNS under pathological conditions (72–77). Thus, we hypothesize that in addition to fungal factors mediating microglia-specific phagocytosis evasion, differences in immunophenotypes and phagocytic receptor expression between microglia and peripheral macrophages are contributing to the defect seen in recognition and engulfment by C20 cells. Although more work is needed to elucidate the differences between these phagocytes, specifically regarding cryptococcal infection, our data is consistent with both fungal and host factors mediating this phenotype.

Consistent with a fungal contribution, we identified four mutants with increased internalization by C20 cells relative to WT. We sought to look for commonalities between these mutants that could explain their differential recognition and engulfment by microglia. We first looked at capsule production and found that none of the mutants had altered capsule production compared to WT, except for *hpi1*Δ, which had a significantly larger capsule radius. However, since the capsule is antiphagocytic due to its ability to hide the pathogen-associated molecular patterns (PAMPs) on the fungal cell surface, is unlikely that an increase in capsule size compared to WT is a contributing factor to this mutant’s enhanced engulfment by C20 cells. However, when we looked at cell body size, overall, the mutants are smaller. In host-like media, we found that all mutants, except for *hir1*Δ, had significantly lower cell body size, and in nutrient-rich media all mutants exhibited significantly lower cell body sizes compared to WT. This phenotype could directly contribute to the ability of these mutants to be engulfed by microglia, as it has been shown that smaller *C. neoformans* cells have increased association with liver macrophages (78). We also looked at the exposure of cryptococcal cell wall components of these mutants compared to WT and found that all mutants had significantly higher intensity of chitin (CFW) and chitosan (EoY) on their cell walls compared to WT. While we were not surprised to find that these mutants have alterations in their cell wall, we were surprised to find a unanimous cell wall phenotype for all the mutants with enhanced engulfment. This could indicate that microglia have increased expression of receptors for chitin and chitosan, or for another PAMP exposed in all of these mutants. We also looked at the ability of these mutants to produce virulence factors such as melanin and urease. We saw that, relative to WT, *hir1*Δ had significantly less shed melanin concomitant with melanin accumulation on the cell wall, and this could be a consequence of cell wall alterations that could potentially enhance recognition and engulfment by microglia. None of the other mutants had defects in melanin and all of them had normal urease production. Lastly, we tested the ability of these mutants to grow under a variety of stressors and found that *hir1*Δ has a significant growth defect at high temperature compared to WT, and it, along with *rim101*Δ, exhibits increased sensitivity to oxidative stress. Thus, the only commonality found was the overall smaller size of these mutants, and the increased exposure of chitin and chitosan.

Lastly, we studied the maturation of WT cryptococcal phagosomes in C20 cells and primary human microglia. Relatively little is known about phagosome maturation in microglia after *C. neoformans* engulfment, despite several studies published about cryptococcal manipulation of phagosome maturation and acidification in peripheral macrophages. We find that WT *C. neoformans* manipulates phagosome maturation in both C20 cells and primary human microglia by retaining EEA1 in and delaying vATPase and LAMP1 recruitment to its phagosomal membrane. Since EEA1 is an early endosomal marker, it still remains unclear whether vATPase and LAMP1 association is perturbed in cryptococcal phagosomes due to delayed EEA1 association and retention after engulfment. Notably, it has been shown that peripheral macrophages accumulate LAMP1 in cryptococcal-containing phagosomes much faster than microglia (16). Specifically, 60-70% of cryptococcal phagosomes are positive for LAMP1 just 30 minutes after coculture with BMDMs (16), compared to the 3 hours it takes for cryptococcal phagosomes to reach this association of LAMP1 after engulfment by microglia. While we already show that microglia respond differently to *C. neoformans* than macrophages, it is important to note that their findings resulted from the use of murine macrophages and antibody-opsonized H99. However, while more direct comparisons need to be done regarding microglia vs macrophage cryptococcal phagosome maturation, we believe that microglia have significant delays in recognition and phagosome maturation compared to other macrophages. Furthermore, while ∼70% of cryptococcal phagosomes become positive for LAMP1 after 3 hours post-coculture with C20 cells and primary human microglia, this does not correlate to fungicidal activity within C20 cells. We assessed phagosome maturation and fungicidal activity of C20 cells against cryptococcal mutants known to have altered intracellular, or *in-vivo*, survival and found that although all mutants have increased association of LAMP1, some were still growing inside microglia. This indicates that alterations in the expression of PAMPs in the cryptococcal cell wall can have drastic effects not only on recognition and engulfment, but also on phagosome maturation. Although studies have shown that both pathogens and host immune cells alter phagosome maturation based on PAMP exposure, and subsequent pattern recognition receptor engagement (79), this is surprising for the cryptococcal field, where typically mutants that cannot alter phagosomal maturation are defective in intracellular growth (21). Thus, the maturation of the cryptococcal phagosome needs to be studied more in-depth to evaluate phagosome acidification and phagolysosome function (i.e., cathepsin activity) in microglia. It is clear that altered PAMP exposure and phagosome maturation does not necessarily correlate with enhanced fungicidal activity by C20 cells.

While many studies have looked at the accumulation of damage on cryptococcal phagosomes in peripheral macrophages, this had not been tested directly in microglia. Notably, in THP-1 cells only 5 to 10% of cryptococcal phagosomes are positive for Gal-3 after 24 hours (21). This is in direct contrast to our results that show almost 60% of phagosomes are positive for Gal-3 staining in C20 cells after 24 hours. While these findings cannot be directly compared, this further supports our hypothesis that microglia are ineffective against *C. neoformans* to an exacerbated extent compared to that of other macrophages. Also, we show that the opsonization status of the fungus have a significant effect on the membrane damage accumulation of cryptococcal phagosomes after 24 hours. We believe this is due to the fact that opsonization activates different receptors on the microglial cell surface resulting in different phagosome maturation dynamics in macrophages (80–82). This highlights the importance of studying the roles of opsonization during cryptococcal infection in the brain, where antibodies are not typically found (they cannot cross the BBB and there are no B-cells under steady state conditions) and complement has functions other than opsonization (i.e. synapse pruning).

In conclusion, we demonstrate the use of C20 cells to study cryptococcal-microglia interactions and have found that microglia recognize and respond to cryptococcal infection differently than that of peripheral macrophages. We also found that *C. neoformans* manipulates phagosome maturation in both immortalized and primary human microglia, and this is dependent on capsule and cell wall modifications. Furthermore, we show that mutants with increased internalization by microglia can be identified using the C20 cell line, and enhanced internalization could be attributed to a smaller cell body size and altered cell wall composition. Overall, we believe C20 cells can be used to better define cryptococcal-microglia interactions by allowing future mechanistic studies that can then be validated *in vivo*.

## 4. Methods

### Strains, cell lines, and growth conditions

Strains used were *C. neoformans* serotype A strain KN99α (83), and *S. cerevisiae* strain BY4741 (84) expressing mCherry (85). For some assays, the *C. neoformans* strain was expressing mCherry (86). The *C. neoformans* mutants were obtained from the Madhani deletion collection (38). All fungal strains were maintained at −80°C and grown at 30°C on yeast-peptone-dextrose (YPD) plates.

The human monocytic cell line THP-1 (ATCC TIB-202) was grown in THP-1 complete media (RPMI-1640 with L-glutamine supplemented with 1 mM sodium pyruvate, 0.05% βmercaptoethanol, 10% heat-inactivated fetal bovine serum (FBS), and 1X Pen-Strep solution (100 units/mL penicillin and 100 μg/ml streptomycin)) and passed 2 times per week. The cells were never used after 12 passages. For differentiation into alveolar macrophages, the THP-1 cells were treated as described (41) except we used 0.16 μM phorbol 12-myristate 12-acetate (PMA, Millipore Sigma).

The human microglia cell line C20 (25) was grown in C20 complete media (DMEM-F12 with 10% FBS, 1X Pen-Strep solution (100 units/mL penicillin and 100 μg/mL streptomycin), and 1X N-2 supplement (Gibco #17502048). Cells were supplemented separately with 1 mM fresh dexamethasone. C20 cells were passaged 2-3 times per week and were never used after 12 passages.

The primary human microglia cells were obtained from Celprogen (#37089-01) and grown in Human Microglia Primary Cell Culture Complete Media with Serum (Celprogen #M37089-01S). Cells were passaged 3-4 times per week and were never used after 8 passages.

### Immunofluorescence

C20 cells were seeded at a density of 2.74×10^5^ cells/mL on collagen-coated coverslips (#1.5 round 12 mm glass coverslips, Electron Microscopy Sciences) that were incubated for 1 hour at room temperature (RT) in 50 μg/mL Rat-Tail Collagen IV (Corning) diluted in 0.02 N acetic acid and washed 3X with DPBS. Primary human microglia were seeded at a density of 2.74×10^5^ cells/mL on #1.5 round 12 mm pre-treated German glass coverslips (Electron Microscopy Sciences). The cells were activated overnight with 0.2 μg/mL TNF-α prior to challenge with fungal strains or latex beads. Fungal strains were grown overnight, harvested in the log phase (OD600 of 0.6-0.8), and washed 2X with PBS. Strains not expressing mCherry were incubated with Lucifer Yellow (250 μg/mL) for 30 mins at 30°C with rotation and washed 2X with PBS. The cell counts were collected (Bio-Rad Cell Counter) and fungal cells were either unopsonized (PBS) or opsonized with 40% human serum (Sigma Aldrich) at a ratio of 1×10^5^ cells/μL of serum. Fungal cells were washed 1X in PBS, resuspended in prewarmed basal media (DMEM-F12), and added to the microglia-adhered coverslips at an MOI of 20:1. In place of fungal strains, 5 μm fluorescent latex beads (CD Bioparticles) were washed 1X in PBS and treated as above. At specified timepoints, the wells containing the C20-*Cryptococcus*/beads were washed with DBPS, fixed with 3.7% formaldehyde in PBS for 10 minutes at RT, washed with PBS, and permeabilized with 0.3% saponin in PBS for 10 minutes at RT. Cells were washed with PBS and blocked for 30 minutes in 10% goat serum in PBS. Cells were immunolabeled in 1% goat serum at 4°C overnight with an antibody against Gal-3 (rat; M3/38: sc23938, Santa Cruz Biotechnology; 1:100 concentration), LAMP1 (mouse; Developmental Studies Hybridoma Bank, H4A3; 1:500 concentration), EEA1 (rabbit; Abcam, ab2900; 1:100 concentration), or vATPase (mouse; Santa Cruz Biotechnology, D-11; 1:500 concentration). Coverslips were washed with PBS and a corresponding Alexa Fluor-conjugated goat secondary antibody (Invitrogen) was diluted to 1 μg/mL in PBS with 2 μg/mL of 4’6-diamidino-2-phenylindole (DAPI; Millipore Sigma) and added to cells for 1 hour at RT. Cells were washed with PBS and coverslips were mounted to imaging slides with 5 μL Prolong Diamond Antifade Mountant (Invitrogen) and allowed to cure for at least 24 hours before imaging. Images of fixed cells were acquired using a Zeiss Axio Observer 7, with an Axiocam 506 mono camera and a 100X/1.4 plan-apo oil objective.

### Uptakes assays

C20 cells/primary human microglia were activated with 0.2 μg/mL TNF-α and seeded on glass-bottom 96-well plates (Corning) at a density of 2.74×10^5^ cells/mL and allowed to adhere overnight. THP-1 cells were seeded at a density of 2.74×10^5^ cells/mL and differentiated with 0.16 μM PMA for 48 hours followed by a recovery step of 24 hours without PMA. The microglia/macrophages were challenged with fungi as above. After 3 hours, the cells were washed, fixed, and permeabilized as above. Cells were then stained with 2 μg/mL of DAPI and 2.5 μg/mL of CellMask Deep Red (Invitrogen) for 10 minutes at RT. Cells were washed with PBS and stored at 4°C in 1 mM NaN_3_ until imaging. The plates were imaged on a Zeiss Axio Observer Microscope with an automatic stage. Each well was imaged using a 3×3 grid set up and resulting images were analyzed using a Cell Profiler (87) pipeline to determine the Phagocytic Index (PI) values.

For heat-killed fungi, *C. neoformans* was harvested in log phase, washed 2X with PBS, and incubated either at 75°C (heat-killed) or RT (alive) for 30 minutes. Cells were washed 1X in PBS and cell counts were attained. For conditioned media, overnight fungal cultures were diluted to 1×10^5^ cells/mL in DMEM/F-12 and incubated for 72 hours at 37°C 5% CO_2_ in a T75 flask (VWR). Cells were removed by centrifugation and the supernatant was filtered through a 0.22 μm filter. The conditioned media was used in place of DMEM-F-12 when preparing the MOI suspension.

### Killing assays

C20 cells were activated with 0.2 μg/mL TNF-α and seeded on 24-well plates (1 per timepoint) at a density of 3.34×10^5^ cells/mL and allowed to adhere overnight. C20 cells were challenged with fungi as above. 3 hours after challenge, plates of all timepoints were washed 2X with DPBS to remove extracellular fungi. 24- and 48-hour plates were fed with DMEM-F-12 and returned to the incubator for their specified timepoints (the 48-hour plate was washed 2X with DPBS after an additional 24-hour incubation and fed with DMEM-F-12). At the specified timepoints after DPBS washes, 900 μL of killing buffer (0.1% Triton X-100, 1 mM EDTA) was added to each well and the cells were lysed for 5 min, 300 rpm at RT. Lysates were diluted, spread on YPD, and CFUs were counted after 2-day incubation at 30°C.

### Isolation of primary murine microglia

Microglia were isolated from C57BL/6 via immunopanning as described (88, 89). Briefly, 6 μg/mL of goat anti-rat IgG secondary antibody (Invitrogen) is diluted in 50 mM Tris-HCl pH 9.5 and added to a 15 mm cell culture dish to prepare the panning plate. The panning plate is incubated at 37°C 5% CO_2_ for 3 hours. The panning dish is rinsed 3X with panning buffer (0.2% BSA in DPBS) and 1 μg/mL of rat anti-mouse CD11b antibody (Invitrogen; M1/70) diluted in panning buffer is added to the panning plate. The plate is incubated at RT overnight. Mouse pups (4-7 days old) are decapitated, and the brains are removed and rinsed 4X in DPBS. The tissue is homogenized using a razor blade and put on ice. Dissociation buffer (200 U papain (Worthington Biochemical LS003126) dissolved in EBSS (Invitrogen), 1X DNase I (Worthington Biochemical LS002007; 500X stock solution prepared by dissolving 12,500 U of DNase I in 1 mL of EBSS)) is added to the brain homogenate and the dish is placed in a 37°C 5% CO_2_ for 30 minutes. Neutralization buffer (20% FBS in DMEM-F-12) is added to the homogenate, moved to a 50 mL conical, and centrifuged at 500 x g for 15 min at 4°C. The pellet is resuspended in 10 mL of microglia media (ScienCell Microglia Medium #1901) and strained using a 70 μm cell strainer. The strained cell mixture is centrifuged at 500 x g for 10 min at 4°C. The pellet is resuspended in panning buffer and added to the panning plate (which has been washed 3X with DPBS). The panning plate is incubated at RT for 20 minutes. The panning plate is rinsed 10X with DPBS. 0.25% trypsin (Corning) is added to the panning plate and left to incubate for 10 minutes at 37°C 5% CO_2_. The panning plate is washed 2X with DPBS, ice-cold microglia medium is added, and the plate is incubated for 2 minutes on ice. A serological pipet is used to dislodge the cells from the panning plate. Cell suspension is collected and centrifuged at 500 x g for 15 min at 4°C. The pellet is resuspended in 500 μL of microglia medium and the cells are counted. 100 μL of a 2×10^5^ cell/mL suspension is aliquoted to a glass-bottom 96-well plate. The uptake assay is performed as above after 14 days.

### Capsule induction

Fungal strains were grown overnight in YPD at 30°C, 250 rpm. The cells were washed once each with PBS and DMEM. 1×10^6^ cells/mL were added to 24-well tissue culture plates. The plates were incubated for 24 hours at 37°C 5% CO_2_. The cell suspension was collected, washed once with PBS, and resuspended in 50 μL PBS. 6 μL of cell resuspension was added to 4 μL of India Ink and samples were visualized on a 100X objective of an inverted Zeiss microscope. Images were analyzed with ImageJ software for capsule radius. Cell size measurements (DMEM condition) were also taken via ImageJ software.

### Cell wall staining

Fungal strains were grown overnight in YPD at 30°C, 250 rpm. The cells were washed twice with PBS. 1×10^7^ cells/mL were stained with 50 μg/mL Alexa Fluor 488-conjugated concanavalin A (ConA) and 100 μg/mL Calcofluor White (CFW) in PBS. For eosin Y (EoY), cells were washed one additional time in McIlvaine’s buffer pH 6.0 and stained with 500 μg/mL EoY in McIlvaine’s. Cells were stained for 30 min at room temperature with gentle shaking. Cells were washed three times with PBS and imaged at 100X using an inverted Zeiss microscope. Fluorescence of ConA, CFW, and EoY were obtained at 488 nm, 405 nm, and 558 nm, respectively. MFI was analyzed using a Cell Profiler (87) pipeline. Cell size measurements (YPD condition) were also taken via ImageJ software.

### Melanin and urease production

Solid and liquid melanin-inducing media were prepared as described (90). Christensen Urea Agar Base was purchased and prepared as directed (Sigma). For solid medium (melanin and urease), fungal strains were grown overnight in YPD at 30°C, 250 rpm, and harvested in the log phase (OD_600_ 0.6-0.8). Cells were washed twice with PBS and the OD_600_ of each strain was normalized to 0.25. 5 μL of each normalized strain was spotted on the solid medium and plates were incubated at 30°C for 72 hours.

For liquid medium (melanin), fungal strains were grown overnight in YPD at 30°C, 250 rpm, and harvested in the log phase (OD_600_ 0.6-0.8). Cells were washed twice with PBS and the OD_600_ of each strain was normalized to 0.25 in a 5 mL culture. Cultures were incubated at 30°C, 250 rpm for 18 hours. The cultures were centrifuged at 3,000 x g for 5 min. The OD_475_ of the supernatant for each strain was measured.

### Stress plates

Fungal strains were grown overnight in YPD at 30°C, 250 rpm, and harvested in the log phase (OD_600_ 0.6-0.8). Cells were washed twice with PBS and diluted to 1×10^7^ cells/mL. 10-fold serial dilutions were performed and 5 of each dilution was spotted onto YPD, RPMI, and YPD with 2 mM H_2_O_2_. YPD plates were incubated at 30°C, 37°C, and 37°C 5% CO_2_ for 48 hours. RPMI plates were incubated at 37°C 5% CO_2_ for 48 hours. 2 mM H_2_O_2_ plates were incubated at 30°C for 48 hours.

## Acknowledgments

Work in the Santiago-Tirado lab is supported by grants from the NIH (R21AI171742 and R01AI177875) and from institutional funds (University of Notre Dame). We acknowledge financial support from the Berthiaume Institute for Precision Health to RLR. We are thankful to Katrina Adams for her support with murine microglia isolation.

## References

1. Hawksworth DL, Lucking R. 2017. Fungal Diversity Revisited: 2.2 to 3.8 Million Species. Microbiol Spectr 5.

2. Kohler JR, Casadevall A, Perfect J. 2014. The spectrum of fungi that infects humans. Cold Spring Harb Perspect Med 5:a019273.

3. Denning DW. 2024. Global incidence and mortality of severe fungal disease. Lancet Infect Dis doi:10.1016/S1473-3099(23)00692-8.

4. Briner SL, Doering TL. 2025. Cryptococcus. Curr Biol 35:R518–R522.

5. Rajasingham R, Govender NP, Jordan A, Loyse A, Shroufi A, Denning DW, Meya DB, Chiller TM, Boulware DR. 2022. The global burden of HIV-associated cryptococcal infection in adults in 2020: a modelling analysis. Lancet Infect Dis doi:10.1016/S1473-3099(22)00499-6.

6. Gaylord EA, Choy HL, Doering TL. 2020. Dangerous Liaisons: Interactions of *Cryptococcus neoformans* with Host Phagocytes. Pathogens 9.

7. Van Hove H, De Feo D, Greter M, Becher B. 2025. Central Nervous System Macrophages in Health and Disease. Annu Rev Immunol doi:10.1146/annurev-immunol-082423-041334.

8. Figarella K, Uzcategui NL, Mogk S, Wild K, Fallier-Becker P, Neher JJ, Duszenko M. 2018. Morphological changes, nitric oxide production, and phagocytosis are triggered in vitro in microglia by bloodstream forms of Trypanosoma brucei. Sci Rep 8:15002.

9. Wu Y, Du S, Bimler LH, Mauk KE, Lortal L, Kichik N, Griffiths JS, Osicka R, Song L, Polsky K, Kasper L, Sebo P, Weatherhead J, Knight JM, Kheradmand F, Zheng H, Richardson JP, Hube B, Naglik JR, Corry DB. 2023. Toll-like receptor 4 and CD11b expressed on microglia coordinate eradication of Candida albicans cerebral mycosis. Cell Rep 42:113240.

10. Neal LM, Xing E, Xu J, Kolbe JL, Osterholzer JJ, Segal BM, Williamson PR, Olszewski MA. 2017. CD4(+) T Cells Orchestrate Lethal Immune Pathology despite Fungal Clearance during Cryptococcus neoformans Meningoencephalitis. mBio 8.

11. Zhou Q, Gault RA, Kozel TR, Murphy WJ. 2007. Protection from direct cerebral cryptococcus infection by interferon-gamma-dependent activation of microglial cells. J Immunol 178:5753–61.

12. Lee SC, Kress Y, Dickson DW, Casadevall A. 1995. Human microglia mediate anti-Cryptococcus neoformans activity in the presence of specific antibody. J Neuroimmunol 62:43–52.

13. Lee SC, Casadevall A, Dickson DW. 1996. Immunohistochemical localization of capsular polysaccharide antigen in the central nervous system cells in cryptococcal meningoencephalitis. Am J Pathol 148:1267–74.

14. Mohamed SH, Fu MS, Hain S, Alselami A, Vanhoffelen E, Li Y, Bojang E, Lukande R, Ballou ER, May RC, Ding C, Velde GV, Drummond RA. 2023. Microglia are not protective against cryptococcal meningitis. Nat Commun 14:7202.

15. Lee SC, Kress Y, Zhao ML, Dickson DW, Casadevall A. 1995. Cryptococcus neoformans survive and replicate in human microglia. Lab Invest 73:871–9.

16. Alvarez M, Casadevall A. 2006. Phagosome extrusion and host-cell survival after *Cryptococcus neoformans* phagocytosis by macrophages. Curr Biol 16:2161–5.

17. Davis MJ, Eastman AJ, Qiu Y, Gregorka B, Kozel TR, Osterholzer JJ, Curtis JL, Swanson JA, Olszewski MA. 2015. *Cryptococcus neoformans*-induced macrophage lysosome damage crucially contributes to fungal virulence. J Immunol 194:2219–31.

18. De Leon-Rodriguez CM, Fu MS, Corbali MO, Cordero RJB, Casadevall A. 2018. The Capsule of *Cryptococcus neoformans* Modulates Phagosomal pH through Its Acid-Base Properties. mSphere 3.

19. De Leon-Rodriguez CM, Rossi DCP, Fu MS, Dragotakes Q, Coelho C, Guerrero Ros I, Caballero B, Nolan SJ, Casadevall A. 2018. The Outcome of the Cryptococcus neoformans-Macrophage Interaction Depends on Phagolysosomal Membrane Integrity. J Immunol 201:583–603.

20. Levitz SM, Nong SH, Seetoo KF, Harrison TS, Speizer RA, Simons ER. 1999. *Cryptococcus neoformans* resides in an acidic phagolysosome of human macrophages. Infect Immun 67:885–90.

21. Santiago-Burgos EJ, Stuckey PV, Santiago-Tirado FH. 2022. Real-time visualization of phagosomal pH manipulation by *Cryptococcus neoformans* in an immune signal-dependent way. Front Cell Infect Microbiol 12:967486.

22. Smith LM, Dixon EF, May RC. 2015. The fungal pathogen *Cryptococcus neoformans* manipulates macrophage phagosome maturation. Cell Microbiol 17:702–13.

23. Tucker SC, Casadevall A. 2002. Replication of *Cryptococcus neoformans* in macrophages is accompanied by phagosomal permeabilization and accumulation of vesicles containing polysaccharide in the cytoplasm. Proc Natl Acad Sci U S A 99:3165–70.

24. Alvarez-Carbonell D, Ye F, Ramanath N, Dobrowolski C, Karn J. 2019. The Glucocorticoid Receptor Is a Critical Regulator of HIV Latency in Human Microglial Cells. J Neuroimmune Pharmacol 14:94–109.

25. Garcia-Mesa Y, Jay TR, Checkley MA, Luttge B, Dobrowolski C, Valadkhan S, Landreth GE, Karn J, Alvarez-Carbonell D. 2017. Immortalization of primary microglia: a new platform to study HIV regulation in the central nervous system. J Neurovirol 23:47–66.

26. Milenkovic VM, Slim D, Bader S, Koch V, Heinl ES, Alvarez-Carbonell D, Nothdurfter C, Rupprecht R, Wetzel CH. 2019. CRISPR-Cas9 Mediated TSPO Gene Knockout alters Respiration and Cellular Metabolism in Human Primary Microglia Cells. Int J Mol Sci 20.

27. Pozzo ED, Tremolanti C, Costa B, Giacomelli C, Milenkovic VM, Bader S, Wetzel CH, Rupprecht R, Taliani S, Settimo FD, Martini C. 2019. Microglial Pro-Inflammatory and Anti-Inflammatory Phenotypes Are Modulated by Translocator Protein Activation. Int J Mol Sci 20.

28. Randall LD, Daniel JB, Kelly M, Gary WC, Subhas D. 2018. Interleukin-1β-induced inflammatory signaling in C20 human microglial cells. Neuroimmunology and Neuroinflammation 5:50.

29. Campuzano A, Wormley FL. 2018. Innate Immunity against *Cryptococcus*, from Recognition to Elimination. J Fungi (Basel) 4.

30. Hole C, Wormley FL, Jr. 2016. Innate host defenses against Cryptococcus neoformans. J Microbiol 54:202–11.

31. Levitz SM. 2010. Innate recognition of fungal cell walls. PLoS Pathog 6:e1000758.

32. Colombo AC, Rella A, Normile T, Joffe LS, Tavares PM, de SAGR, Frases S, Orner EP, Farnoud AM, Fries BC, Sheridan B, Nimrichter L, Rodrigues ML, Del Poeta M. 2019. Cryptococcus neoformans Glucuronoxylomannan and Sterylglucoside Are Required for Host Protection in an Animal Vaccination Model. mBio 10.

33. Enriquez V, Munzen ME, Porras LM, Charles-Nino CL, Yu F, Alvina K, Ramos RL, Dores MR, Giusti-Rodriguez P, Martinez LR. 2025. Active Cryptococcus neoformans glucuronoxylomannan production prevents elimination of cryptococcal CNS infection in vivo. J Neuroinflammation 22:61.

34. Lee HH, Carmichael DJ, Ribeiro V, Parisi DN, Munzen ME, Charles-Nino CL, Hamed MF, Kaur E, Mishra A, Patel J, Rooklin RB, Sher A, Carrillo-Sepulveda MA, Eugenin EA, Dores MR, Martinez LR. 2023. Glucuronoxylomannan intranasal challenge prior to Cryptococcus neoformans pulmonary infection enhances cerebral cryptococcosis in rodents. PLoS Pathog 19:e1010941.

35. Chun CD, Brown JCS, Madhani HD. 2011. A major role for capsule-independent phagocytosis-inhibitory mechanisms in mammalian infection by Cryptococcus neoformans. Cell Host Microbe 9:243–251.

36. Chun CD, Madhani HD. 2010. Ctr2 links copper homeostasis to polysaccharide capsule formation and phagocytosis inhibition in the human fungal pathogen Cryptococcus neoformans. PLoS One 5.

37. Gaylord EA, Choy HL, Chen G, Briner SL, Doering TL. 2024. Sac1 links phosphoinositide turnover to cryptococcal virulence. mBio 15:e0149624.

38. Liu OW, Chun CD, Chow ED, Chen C, Madhani HD, Noble SM. 2008. Systematic genetic analysis of virulence in the human fungal pathogen Cryptococcus neoformans. Cell 135:174–88.

39. Luberto C, Martinez-Marino B, Taraskiewicz D, Bolanos B, Chitano P, Toffaletti DL, Cox GM, Perfect JR, Hannun YA, Balish E, Del Poeta M. 2003. Identification of App1 as a regulator of phagocytosis and virulence of Cryptococcus neoformans. J Clin Invest 112:1080–94.

40. Price MS, Nichols CB, Alspaugh JA. 2008. The Cryptococcus neoformans Rho-GDP dissociation inhibitor mediates intracellular survival and virulence. Infect Immun 76:5729–37.

41. Santiago-Tirado FH, Peng T, Yang M, Hang HC, Doering TL. 2015. A Single Protein S-acyl Transferase Acts through Diverse Substrates to Determine Cryptococcal Morphology, Stress Tolerance, and Pathogenic Outcome. PLoS Pathog 11:e1004908.

42. Winski CJ, Qian Y, Mobashery S, Santiago-Tirado FH. 2022. An Atypical ABC Transporter Is Involved in Antifungal Resistance and Host Interactions in the Pathogenic Fungus *Cryptococcus neoformans*. mBio 13:e0153922.

43. Bojarczuk A, Miller KA, Hotham R, Lewis A, Ogryzko NV, Kamuyango AA, Frost H, Gibson RH, Stillman E, May RC, Renshaw SA, Johnston SA. 2016. Cryptococcus neoformans Intracellular Proliferation and Capsule Size Determines Early Macrophage Control of Infection. Sci Rep 6:21489.

44. Chang YC, Kwon-Chung KJ. 1994. Complementation of a capsule-deficient mutation of Cryptococcus neoformans restores its virulence. Mol Cell Biol 14:4912–9.

45. Mylonakis E, Moreno R, El Khoury JB, Idnurm A, Heitman J, Calderwood SB, Ausubel FM, Diener A. 2005. Galleria mellonella as a model system to study Cryptococcus neoformans pathogenesis. Infect Immun 73:3842–50.

46. Winski CJ, Stuckey PV, Marrufo AM, Agyei G, Ross RL, Urmi T, Chapman S, Santiago-Tirado FH. 2025. Lack of an atypical PDR transporter generates an immunogenic Cryptococcus neoformans strain that drives a dysregulated and lethal immune response in murine lungs. mBio doi:10.1128/mbio.01321-25:e0132125.

47. Kraus PR, Fox DS, Cox GM, Heitman J. 2003. The Cryptococcus neoformans MAP kinase Mpk1 regulates cell integrity in response to antifungal drugs and loss of calcineurin function. Mol Microbiol 48:1377–87.

48. Hu G, Caza M, Cadieux B, Bakkeren E, Do E, Jung WH, Kronstad JW. 2015. The endosomal sorting complex required for transport machinery influences haem uptake and capsule elaboration in Cryptococcus neoformans. Mol Microbiol 96:973–92.

49. Aits S, Kricker J, Liu B, Ellegaard AM, Hamalisto S, Tvingsholm S, Corcelle-Termeau E, Hogh S, Farkas T, Holm Jonassen A, Gromova I, Mortensen M, Jaattela M. 2015. Sensitive detection of lysosomal membrane permeabilization by lysosomal galectin puncta assay. Autophagy 11:1408–24.

50. Paz I, Sachse M, Dupont N, Mounier J, Cederfur C, Enninga J, Leffler H, Poirier F, Prevost MC, Lafont F, Sansonetti P. 2010. Galectin-3, a marker for vacuole lysis by invasive pathogens. Cell Microbiol 12:530–44.

51. Francis VI, Liddle C, Camacho E, Kulkarni M, Junior SRS, Harvey JA, Ballou ER, Thomson DD, Brown GD, Hardwick JM, Casadevall A, Witton J, Coelho C. 2024. Cryptococcus neoformans rapidly invades the murine brain by sequential breaching of airway and endothelial tissues barriers, followed by engulfment by microglia. mBio 15:e0307823.

52. Nielson JA, Davis JM. 2023. Roles for Microglia in Cryptococcal Brain Dissemination in the Zebrafish Larva. Microbiol Spectr 11:e0431522.

53. Blasi E, Barluzzi R, Mazzolla R, Tancini B, Saleppico S, Puliti M, Pitzurra L, Bistoni F. 1995. Role of nitric oxide and melanogenesis in the accomplishment of anticryptococcal activity by the BV-2 microglial cell line. J Neuroimmunol 58:111–6.

54. Hamed MF, Araujo GRS, Munzen ME, Reguera-Gomez M, Epstein C, Lee HH, Frases S, Martinez LR. 2023. Phospholipase B Is Critical for Cryptococcus neoformans Survival in the Central Nervous System. mBio 14:e0264022.

55. Redlich S, Ribes S, Schutze S, Eiffert H, Nau R. 2013. Toll-like receptor stimulation increases phagocytosis of Cryptococcus neoformans by microglial cells. J Neuroinflammation 10:71.

56. Abels ER, Nieland L, Hickman S, Broekman MLD, El Khoury J, Maas SLN. 2021. Comparative Analysis Identifies Similarities between the Human and Murine Microglial Sensomes. Int J Mol Sci 22.

57. Hoos MD, Vitek MP, Ridnour LA, Wilson J, Jansen M, Everhart A, Wink DA, Colton CA. 2014. The impact of human and mouse differences in NOS2 gene expression on the brain’s redox and immune environment. Mol Neurodegener 9:50.

58. Horvath RJ, Nutile-McMenemy N, Alkaitis MS, Deleo JA. 2008. Differential migration, LPS-induced cytokine, chemokine, and NO expression in immortalized BV-2 and HAPI cell lines and primary microglial cultures. J Neurochem 107:557–69.

59. Lee SC, Liu W, Dickson DW, Brosnan CF, Berman JW. 1993. Cytokine production by human fetal microglia and astrocytes. Differential induction by lipopolysaccharide and IL-1 beta. J Immunol 150:2659–67.

60. Nieman AN, Li G, Zahn NM, Mian MY, Mikulsky BN, Hoffman DA, Wilcox TM, Kehoe AS, Luecke IW, Poe MM, Alvarez-Carbonell D, Cook JM, Stafford DC, Arnold LA. 2020. Targeting Nitric Oxide Production in Microglia with Novel Imidazodiazepines for Nonsedative Pain Treatment. ACS Chem Neurosci 11:2019–2030.

61. Peterson PK, Hu S, Anderson WR, Chao CC. 1994. Nitric oxide production and neurotoxicity mediated by activated microglia from human versus mouse brain. J Infect Dis 170:457–60.

62. Kozel TR, Gotschlich EC. 1982. The capsule of cryptococcus neoformans passively inhibits phagocytosis of the yeast by macrophages. J Immunol 129:1675–80.

63. Small JM, Mitchell TG. 1989. Strain variation in antiphagocytic activity of capsular polysaccharides from Cryptococcus neoformans serotype A. Infect Immun 57:3751–6.

64. Srikanta D, Yang M, Williams M, Doering TL. 2011. A sensitive high-throughput assay for evaluating host-pathogen interactions in *Cryptococcus neoformans* infection. PLoS One 6:e22773.

65. Jain N, Cordero RJ, Casadevall A, Fries BC. 2013. Allergen1 regulates polysaccharide structure in Cryptococcus neoformans. Mol Microbiol 88:713–27.

66. Stano P, Williams V, Villani M, Cymbalyuk ES, Qureshi A, Huang Y, Morace G, Luberto C, Tomlinson S, Del Poeta M. 2009. App1: an antiphagocytic protein that binds to complement receptors 3 and 2. J Immunol 182:84–91.

67. Gallerand A, Han J, Ivanov S, Randolph GJ. 2024. Mouse and human macrophages and their roles in cardiovascular health and disease. Nat Cardiovasc Res 3:1424–1437.

68. Mestas J, Hughes CC. 2004. Of mice and not men: differences between mouse and human immunology. J Immunol 172:2731–8.

69. Saas P, Chague C, Maraux M, Cherrier T. 2020. Toward the Characterization of Human Pro-Resolving Macrophages? Front Immunol 11:593300.

70. Saito Y, Fujiwara Y, Yamaguchi YL, Tanaka SS, Miura K, Hizukuri Y, Yamashiro K, Hayashi Y, Nakashima Y, Komohara Y. 2025. Rodent monocyte-derived macrophages do not express CD163: Comparative analysis using macrophages from living boreoeutherians. Dev Dyn doi:10.1002/dvdy.70036.

71. Vijayan V, Pradhan P, Braud L, Fuchs HR, Gueler F, Motterlini R, Foresti R, Immenschuh S. 2019. Human and murine macrophages exhibit differential metabolic responses to lipopolysaccharide - A divergent role for glycolysis. Redox Biol 22:101147.

72. Cuadros MA, Sepulveda MR, Martin-Oliva D, Marin-Teva JL, Neubrand VE. 2022. Microglia and Microglia-Like Cells: Similar but Different. Front Cell Neurosci 16:816439.

73. DePaula-Silva AB, Gorbea C, Doty DJ, Libbey JE, Sanchez JMS, Hanak TJ, Cazalla D, Fujinami RS. 2019. Differential transcriptional profiles identify microglial- and macrophage-specific gene markers expressed during virus-induced neuroinflammation. J Neuroinflammation 16:152.

74. Li Q, Barres BA. 2018. Microglia and macrophages in brain homeostasis and disease. Nat Rev Immunol 18:225–242.

75. Norris GT, Kipnis J. 2019. Immune cells and CNS physiology: Microglia and beyond. J Exp Med 216:60–70.

76. Prinz M, Masuda T, Wheeler MA, Quintana FJ. 2021. Microglia and Central Nervous System-Associated Macrophages-From Origin to Disease Modulation. Annu Rev Immunol 39:251–277.

77. Yin J, Valin KL, Dixon ML, Leavenworth JW. 2017. The Role of Microglia and Macrophages in CNS Homeostasis, Autoimmunity, and Cancer. J Immunol Res 2017:5150678.

78. Denham ST, Brammer B, Chung KY, Wambaugh MA, Bednarek JM, Guo L, Moreau CT, Brown JCS. 2022. A dissemination-prone morphotype enhances extrapulmonary organ entry by *Cryptococcus neoformans*. Cell Host Microbe 30:1382–1400 e8.

79. Pauwels AM, Trost M, Beyaert R, Hoffmann E. 2017. Patterns, Receptors, and Signals: Regulation of Phagosome Maturation. Trends Immunol 38:407–422.

80. Erwig LP, McPhilips KA, Wynes MW, Ivetic A, Ridley AJ, Henson PM. 2006. Differential regulation of phagosome maturation in macrophages and dendritic cells mediated by Rho GTPases and ezrin-radixin-moesin (ERM) proteins. Proc Natl Acad Sci U S A 103:12825–30.

81. Moretti J, Blander JM. 2014. Insights into phagocytosis-coupled activation of pattern recognition receptors and inflammasomes. Curr Opin Immunol 26:100–10.

82. Rodriguez-Gomez JA, Kavanagh E, Engskog-Vlachos P, Engskog MKR, Herrera AJ, Espinosa-Oliva AM, Joseph B, Hajji N, Venero JL, Burguillos MA. 2020. Microglia: Agents of the CNS Pro-Inflammatory Response. Cells 9.

83. Nielsen K, Cox GM, Wang P, Toffaletti DL, Perfect JR, Heitman J. 2003. Sexual cycle of *Cryptococcus neoformans* var. grubii and virulence of congenic a and alpha isolates. Infect Immun 71:4831–41.

84. Winzeler EA, Shoemaker DD, Astromoff A, Liang H, Anderson K, Andre B, Bangham R, Benito R, Boeke JD, Bussey H, Chu AM, Connelly C, Davis K, Dietrich F, Dow SW, El Bakkoury M, Foury F, Friend SH, Gentalen E, Giaever G, Hegemann JH, Jones T, Laub M, Liao H, Liebundguth N, Lockhart DJ, Lucau-Danila A, Lussier M, M’Rabet N, Menard P, Mittmann M, Pai C, Rebischung C, Revuelta JL, Riles L, Roberts CJ, Ross-MacDonald P, Scherens B, Snyder M, Sookhai-Mahadeo S, Storms RK, Veronneau S, Voet M, Volckaert G, Ward TR, Wysocki R, Yen GS, Yu K, Zimmermann K, Philippsen P, et al. 1999. Functional characterization of the S. cerevisiae genome by gene deletion and parallel analysis. Science 285:901–6.

85. Santiago-Tirado FH, Onken MD, Cooper JA, Klein RS, Doering TL. 2017. Trojan Horse Transit Contributes to Blood-Brain Barrier Crossing of a Eukaryotic Pathogen. MBio 8.

86. Upadhya R, Lam WC, Maybruck BT, Donlin MJ, Chang AL, Kayode S, Ormerod KL, Fraser JA, Doering TL, Lodge JK. 2017. A fluorogenic *C. neoformans* reporter strain with a robust expression of m-cherry expressed from a safe haven site in the genome. Fungal Genet Biol 108:13–25.

87. Carpenter AE, Jones TR, Lamprecht MR, Clarke C, Kang IH, Friman O, Guertin DA, Chang JH, Lindquist RA, Moffat J, Golland P, Sabatini DM. 2006. CellProfiler: image analysis software for identifying and quantifying cell phenotypes. Genome Biol 7:R100.

88. Bohlen CJ, Bennett FC, Bennett ML. 2019. Isolation and Culture of Microglia. Curr Protoc Immunol 125:e70.

89. Emery B, Dugas JC. 2013. Purification of oligodendrocyte lineage cells from mouse cortices by immunopanning. Cold Spring Harb Protoc 2013:854–68.

90. Nichols CB. 2021. Visualization and Documentation of Capsule and Melanin Production in Cryptococcus neoformans. Curr Protoc 1:e27.

